# Genetically Encoded and Modular SubCellular Organelle Probes (GEM-SCOPe) reveal lysosomal and mitochondrial dysfunction driven by *PRKN* knockout

**DOI:** 10.1101/2024.05.21.594886

**Authors:** Camille Goldman, Tatyana Kareva, Lily Sarrafha, Braxton R. Schuldt, Abhishek Sahasrabudhe, Tim Ahfeldt, Joel W. Blanchard

**Author notes:** 2^nd^ corresponding author Correspondence.

## Abstract

Cellular processes including lysosomal and mitochondrial dysfunction are implicated in the development of many diseases. Quantitative visualization of mitochondria and lysosomes is crucial to understand how these organelles are dysregulated during disease. To address a gap in live-imaging tools, we developed GEM-SCOPe (Genetically Encoded and Modular SubCellular Organelle Probes), a modular toolbox of fluorescent markers designed to inform on localization, distribution, turnover, and oxidative stress of specific organelles. We expressed GEM-SCOPe in differentiated astrocytes and neurons from a human pluripotent stem cell *PRKN-*knockout model of Parkinson’s disease and identified disease-associated changes in proliferation, lysosomal distribution, mitochondrial transport and turnover, and reactive oxygen species. We demonstrate GEM-SCOPe is a powerful panel that provide critical insight into the subcellular mechanisms underlying Parkinson’s disease in human cells. GEM-SCOPe can be expanded upon and applied to a diversity of cellular models to glean an understanding of the mechanisms that promote disease onset and progression.

## Introduction

Parkinson’s Disease (PD) is a common neurodegenerative disease that robs individuals of their motor and cognitive functions, affecting over 10 million people worldwide and with an estimated 90,000 new diagnoses in the United States each year^1^. PD is clinically characterized by bradykinesia, resting tremor, rigidity, and postural instability^2–5^ and pathologically characterized by the loss of dopaminergic neurons in the substantia nigra pars compacta and buildup of intracellular deposits of α-synuclein into insoluble protein aggregates called Lewy Bodies^2,6–8^. From the first pathological descriptions of PD in the mid-20^th^ century, advancements in many fields, including histology, genetics, cell biology, and neurology enabled insight into many aspects of the disease, from determining genetic and environmental risk factors and mapping disease progression to identifying pathways and proteins to target therapeutically. Despite great progress, there is still incomplete understanding of the cellular dysfunction that precedes and contributes to chronic neuronal death.

This limited advancement arises in part due to the absence of tools used to track molecular and cellular changes in real-time throughout disease progression. Studies on human post-mortem brain tissue are largely representative of end-stage disease states, making it difficult to dissect which phenotypes are a primary, causative effect and which phenotypes are a secondary response^9^. Mouse models offer more opportunities to study multiple time points throughout disease progression as motor and cognitive symptoms can be monitored over time. However, targeting specific cell types in mice remains technically challenging and time-consuming. Emerging technologies such as the miniscope allow continuous monitoring of fluorescence-based readouts on a cellular level in live mouse and rat brains^10–12^. However, the type of readout from the miniscope has been restricted to neuronal activity from fluorescent calcium sensors and resolution along the z-axis remains a limiting factor in analyzing deeper cortical tissue.

Regardless of such technologies, current genetic rodent models of PD fail to completely recapitulate human disease and lack key pathological hallmarks. Drug-induced models can recapitulate late-stage disease phenotypes and behaviors, but due to their acute onset, are inadequate to study the gradual accumulation of cellular dysfunction that precedes clinical presentation^13,14^. Advancements in induced pluripotent stem cell (iPSC) differentiation and CRISPR/Cas9 genome editing have led to the emergence of genetic cellular models of PD^15–18^. These models enable researchers to study the effects of genetic and environmental factors on specific human cell populations relevant to PD.

Cellular models of PD provide an exciting opportunity to study subcellular changes and responses to stress or treatment in live cells. Cells can be genetically engineered to express fluorescent proteins that can then be visualized by microscopy in live cells. The applicability of GFP as a reporter gene was immediately appreciated after its identification and isolation in 1962^19^. Since then, hundreds of fluorophores have been developed by introducing mutations to naturally occurring fluorophores isolated from sea anemones and jellyfish to modulate fluorescence intensity, half-life, and excitation/emission spectra^20^. Additional fluorophores have been developed as biosensors, by fluorescing or changing fluorescence emission in the presence of a certain ligand or under specific cellular conditions^20–25^. Cells can express these fluorophores with virtually limitless possibilities with regards to subcellular localization, cell-type specific expression, and temporally controlled expression. Genetically encoded fluorescent proteins can be used to follow a population of cells as they change due to aging, genetics, or environmental stressors and stimuli.

PD has a complex genetic architecture underlying the disease. Over 90% of PD cases are idiopathic; over 90 genetic risk loci have been identified by GWAS, but any of those individual variants confer little risk on their own^26^. The remaining 10% of PD cases can be linked to highly penetrant mutations in a handful of genes. While extremely rare, studying the function of these genes and their impact on cellular processes sheds light on the fundamental pathways that drive PD pathogenesis. Here, we focus on the *PRKN* gene. Autosomal recessive loss of function mutations in *PRKN* are the most frequent known genetic cause of early onset cases of PD, accounting for about 15% of PD cases with onset before the age of 50^27–30^. *PRKN* encodes for the protein PRKN, an E3 ubiquitin ligase associated with mitochondria. PRKN directs the ubiquitination of outer mitochondrial membrane proteins on damaged mitochondria, targeting them for degradation via the autophagy-lysosomal pathway (mitophagy)^31–33^. While the function of PRKN is well understood, it is still unclear how loss of PRKN function leads to the gradual disease progression of early onset PD. It is believed that neurodegeneration can begin over 20 years before clinical onset, with a prolonged prodromal phase associated with several non-motor symptoms. Thus, it is crucial to study models of early disease to understand what cellular disturbances in the prodromal phase culminate in neurodegeneration.

Here, we present GEM-SCOPe (**G**enetically **E**ncoded and **M**odular **S**ub**C**ellular **O**rganelle **P**robes), an expandable panel of genetically encoded reporters to track and quantify the subcellular phenotypes in live cells. We developed a library of constructs that localize specifically to the nucleus, mitochondria, and lysosomes and included fluorophores with a variety of emission spectra or biosensor capabilities. All the lentiviral constructs were developed in the same backbone vector to be modular, enabling any component, be it promotor, localization signal, fluorophore, or antibiotic resistance, to be easily removed or replaced. We validated the fluorophores with live-imaging of human induced pluripotent stem cell (hiPSC)-derived midbrain astrocytes employing existing chemical dyes and chemical perturbations. We then applied GEM-SCOPe to hiPSC-derived astrocytes and neurons from a *PRKN*-knockout model of PD,^15^ to demonstrate the applicability in detecting genetically-mediated effects in disease models. This revealed previously unseen changes in cellular proliferation, lysosomal distribution, mitochondrial motility and turnover, and reactive oxygen species production. Our results demonstrate the widespread utility of GEM-SCOPe to study a variety of translationally important subcellular changes. The modular, combinatorial flexibility of GEM-SCOPe can be adapted to investigate any disease, and, therefore, it is a critical new resource that can be broadly applied in neurodegenerative research and beyond. Results

### Building a modular toolbox for the subcellular localization of genetically encoded fluorophores

We developed GEM-SCOPe for live-cell imaging of subcellular organelle dynamics. We aimed to produce a system that is highly modular to easily customize the expression, localization, fluorescence, and selection for any given experimental need. We started with the FUW backbone^34^, a 3^rd^ generation lentivirus backbone with a ubiquitin C promoter for high, ubiquitous expression of the transgene across cell types^34,35^ and a woodchuck hepatitis virus post-transcriptional regulatory element (WPRE) for improved lentiviral expression^34,36,37^. The designed lentiviral constructs contain four components: promoter, localization sequence, fluorophore, and antibiotic resistance (**Fig. 1A**). The content of these four components can be mixed and matched to generate a library of lentiviral constructs to fit a diverse field of experimental needs. To generate constructs *de novo*, components were combined and inserted into the plasmid via Gibson assembly (**Fig. 1B, top**). Restriction sites were kept between each domain, so each component or group of components could be excised via restriction digest and replaced with a different component, either by restriction site cloning or Gibson assembly (**Fig. 1B, bottom**). Thus, each component of the lentiviral plasmid can be removed or replaced to accommodate features such as cell-type specificity with cell-type specific promoters, alternative subcellular localization, and unique fluorophores.

**Fig. 1:**
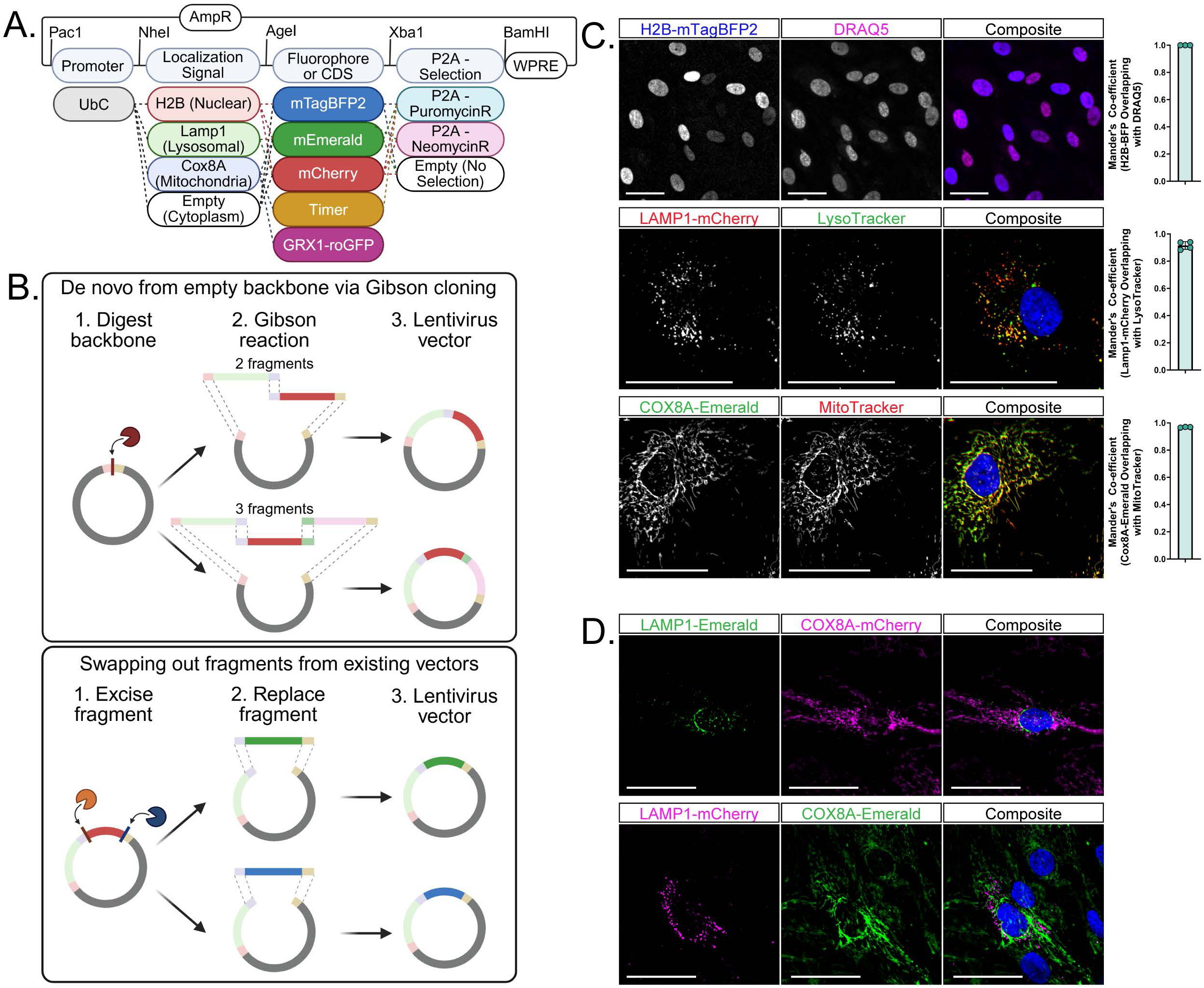
A toolbox of lentiviral plasmids developed and validated for subcellular localization of fluorophores. **(A)** Schematic of lentiviral plasmids highlighting the combinatorial power of the toolbox components. Generated using BioRender **(B)** Schematic demonstrating the cloning strategies used to generate the constructs used in this study as well as any future constructs. Generated using BioRender **(C)** Representative images of subcellular localization validation with available dyes in hiPSC-derived astrocytes and Mander’s co-efficient for overlap of the genetically encoded fluorophore with the dye. Bars represent mean values and error bars represent standard deviation. Top: H2B-mTagBFP2 and DRAQ5 (n = 3; each replicate is an average over 16 images); Middle: LAMP1-mCherry and LysoTracker Deep Red; nuclei stained with Hoechst33342 (n = 4; each replicate is an average over 4 images); Bottom: COX8A-Emerald and MitoTracker Deep Red; nuclei stained with Hoechst33342 (n = 3, each replicate is an average over 36 images). Scale bars = 50 µm **(D)** Representative images of combinatorial applications of GEM-SCOPe. Top: Cell expressing LAMP1-Emerald and COX8A-mCherry. Bottom: cell expression LAMP1-mCherry and COX8A-Emerald. Nuclei were stained with DRAQ5. Scale bars = 50 µm. See also Figure S1 and Table S1.

The FUW backbone has a ubiquitin C promoter, but this can be replaced with lineage or cell-type-specific promoters for selective expression in specific cell types or with drug-inducible promoters to temporally control when the construct is transcribed (**Fig. S1A**). Subcellular localization is determined by including the whole or partial coding sequence of a peptide with a known and specific subcellular localization. This domain can be omitted to produce a fluorophore expressed in the cytoplasm. The fluorophore component of the construct offers the most flexibility, from basic fluorescent proteins to more complex fluorophores that change fluorescent excitation and/or emission wavelength with time or environment. All plasmids generated for GEM-SCOPe have been deposited to Addgene (**Table S1**).

We developed GEM-SCOPe constructs to localize fluorophores to the cytoplasm (no tag), nucleus (H2B-tag), mitochondria (COX8A-tag), and lysosome (LAMP1-tag) (**Fig. 1C**). Fluorophore localization was validated using commercially available live cellular dyes. Nuclear localization was achieved by fusing fluorophores to the C-terminus of histone 2B (H2B). H2B is an integral chromatin protein that is found in all cells and is localized specifically to the nucleus^38,39^. H2B fused to mTagBFP2, a blue fluorophore, (H2B-mTagBFP2) co-localizes with DRAQ5, a far-red live-cell nuclear dye^40^ (**Fig. 1C, top**; Mander’s coefficient > 0.99±0.0001). In addition to the mTagBFP2 construct, we also developed lentiviral plasmids with H2B fused to Emerald (green; H2B-Emerald) and mCherry (red; H2B-mCherry) (**Fig. 1A, S1B**).

To localize fluorophores to the lysosome, we utilized the N-terminal peptide sequence of lysosomal-associated membrane protein 1 (LAMP1), a major component of lysosomal membranes that plays a key role in lysosomal biogenesis and homeostasis. The N-terminus of LAMP1 fused to mCherry (LAMP1-mCherry) co-localizes with LysoTracker (Thermo Scientific) (**Fig. 1C, middle**; Mander’s coefficient = 0.92±0.03). LysoTracker is a fluorophore that is partially protonated at neutral pH and can readily cross membranes; once protonated it can no longer diffuse so the fluorescent signal gets trapped in acidic compartments, including, but not limited to, the lysosome^41,42^. Overexpression of LAMP1 is associated with numerous cancers and cancer metastasis^43–46^. By only using a partial sequence of LAMP1 and not the full peptide sequence, we are able to target the fluorophore specifically to the lysosome without increasing the expression of endogenous LAMP1 (**Fig. S1C**). We also developed lentiviral plasmids with the LAMP1 targeting sequence fused to Emerald (green; LAMP1-Emerald; **Fig. 1A, 1D**).

Finally, mitochondrial localization was achieved using the mitochondrial targeting sequence of COX8A. COX8A is a nuclear-encoded subunit of cytochrome-c oxidase (Complex IV) in the electron transport chain^47,48^. The first 25 amino acid residues of the COX8A peptide sequence target the peptide to the mitochondrial inner membrane and are widely used to specifically localize proteins and fluorophores to the mitochondria^49–51^. To confirm mitochondrial specificity of COX8A-Emerald we compared localization with MitoTracker (Thermo Scientific), a dye that accumulates in the mitochondria due to the negative membrane potential of the inner mitochondrial membrane^41,52^. The mitochondrial targeting sequence fused to an Emerald fluorophore (COX8A-Emerald) exhibited significant co-localization with MitoTracker (**Fig. 1C, bottom**; Mander’s coefficient = 0.97±0.003). We developed lentiviral plasmids with the COX8A targeting sequence fused to mCherry (red; COX8A-mCherry; **Fig. 1A, 1D**), Timer (green/red; COX8A-Timer; **Fig. 1A**, **Fig. 5**), and Lemon (cyan/yellow; COX8A-Lemon; **Fig. 1A, Fig. S1D, E**).

Once the localization strategies were finalized, we swapped out the fluorophores with different fluorescent proteins that are excited and emitted at alternative wavelengths. Mixing localization sequences and fluorophores with unique emission spectra allows simultaneous imaging of multiple sub-cellular processes in the same cells such as mitochondrial network and lysosomal distribution (**Fig. 1D**). Utilizing a combination of subcellular localization and diverse fluorophores, we developed GEM-SCOPe, a fluorescent, live-cell imaging toolbox of lentiviral constructs that were systematically validated to quantify cellular proliferation, lysosomal distribution, and mitochondrial dynamics in human Parkinson’s disease astrocytes.

### H2B-fused fluorophores offer improvements for efficiently labelling nuclei for multi-day live-cell assays

Quantification of cellular phenotypes by fluorescence microscopy is dependent on normalization to total cell number. This is often accomplished by counting the number of nuclei. A few fluorescent dyes that can cross the cellular and nuclear membranes of live cells and bind specifically to DNA are commercially available. While efficient for short-term and end-point assays, such dyes intercalate in DNA and therefore have cytotoxic and mutagenic effects^53,54^. Furthermore, nuclear dyes do not remain contained within the nuclear compartment, and their signal leaches into the cytoplasm in just a few hours after exposing the cells. The cytotoxicity and loss of precise labeling over time render these dyes less useful for long-term microscopy or cell-tracking experiments. To address these concerns, we developed lentiviral constructs expressing nuclear-localized fluorophores by fusing the nucleus-specific histone protein, H2B, with either mTagBFP2, Emerald, or mCherry (**Fig. 1C, 2A**). Human induced pluripotent stem-cell (hiPSC)-derived astrocytes transfected with H2B-mTagBFP2 were co-stained with DRAQ5, a far-red DNA-stain for live cells. The localization of the two nuclear signals was compared 1 hours and 24 hours after the addition of DRAQ5. Immediately after adding DRAQ5, there was no difference between the DRAQ5 stain and the H2B-mTagBFP2 (2-way ANOVA with Tukey’s HSD; adj-p = 0.35) (**Fig. 2B, C**). Over the experimental period, the DRAQ5 signal spread significantly (19±2% extranuclear; adj-p < 0.0001), while the extranuclear H2B-mTagBFP2 signal did not significantly change (adj-p = 0.95) (**Fig. 2B, C**). Although the human eye can still readily distinguish the nucleus from the extranuclear stain in the DRAQ5 images, image analysis programs that rely on grey values for segmentation often fail to correctly differentiate the true nuclear signal from the extranuclear signal. Thus, lentiviral constructs expressing nuclear-localized fluorophores, such as the demonstrated H2B-mTagBFP2, can provide better alternatives for long-term live-imaging experiments with multiple time points.

**Fig. 2:**
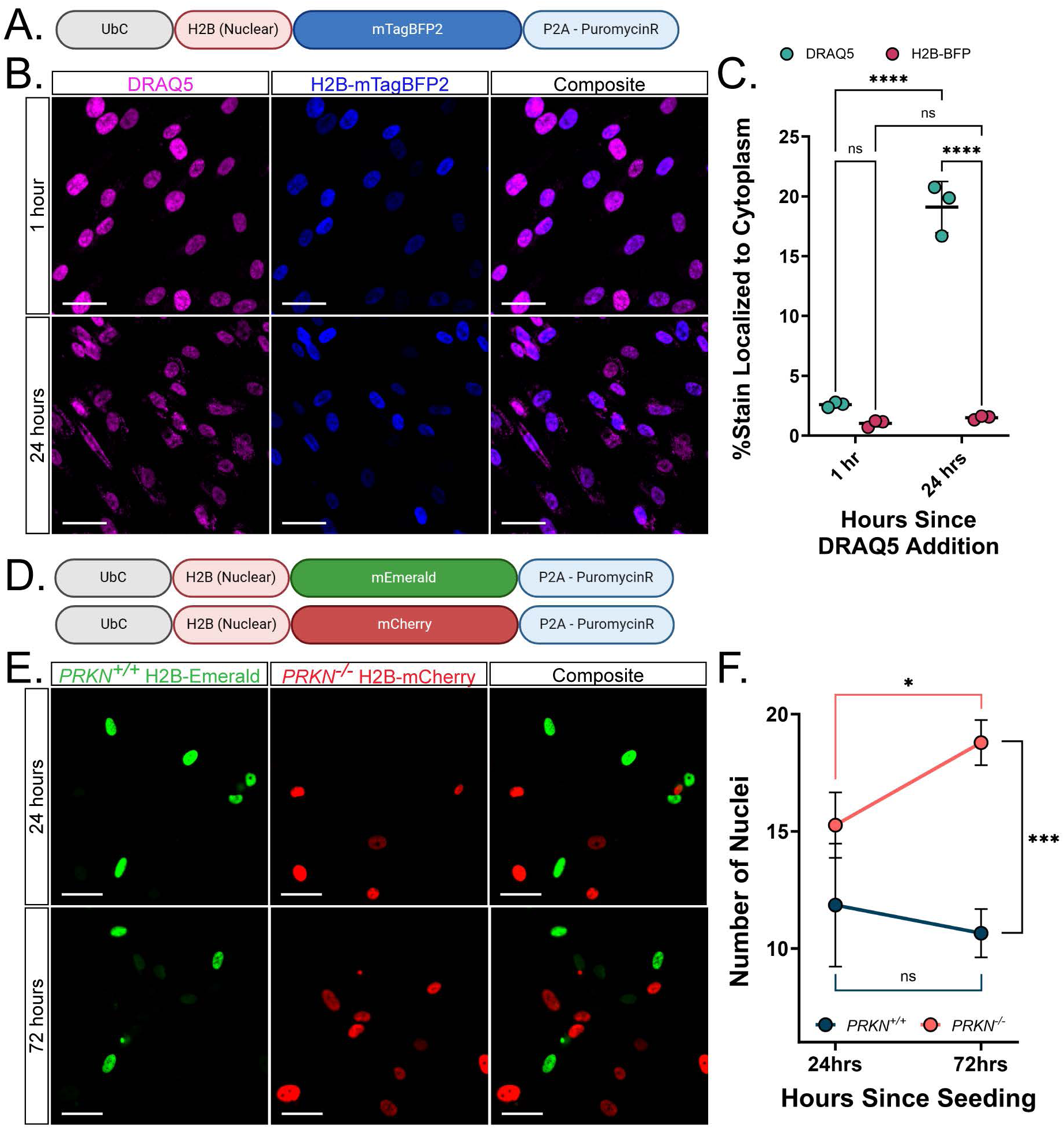
Nucleus-localized fluorophores offer improvements over available stains for multiday imaging. **(A)** Schematic of the lentiviral construct localizing mTagBFP2 to the nucleus with an H2B fusion protein and antibiotic resistance to puromycin used in the following panels **(B)** Representative images of H2B-mTagBFP2 and DRAQ5 nuclear stain 1 hour (top) and 24 hours (bottom) after adding DRAQ5 to the cell cultures. **(C)** Quantification of the percent of stain localized to the nuclear area versus the cytoplasm 1 hour and 24 hours after adding DRAQ5 to cell cultures. The central bar represents the mean and the error bars represent the standard deviation (2-way ANOVA with Tukey’s HSD; n = 3 per time point; each replicate is an average over 36 images) **(D)** Schematic of the lentiviral construct localizing Emerald or mCherry to the nucleus with an H2B fusion protein and antibiotic resistance to puromycin used in the following panels **(E)** Representative images of nuclei from *PRKN^+/+^* astrocytes, labeled with H2B-Emerald, and *PRKN ^-/-^* astrocytes, labeled with H2B-mCherry, cultured together during a proliferation assay. Images were taken 24 hours (top) after seeding and 72 hours (bottom) after seeding **(F)** Quantification of number of *PRKN^+/+^* and *PRKN^-/-^* nuclei per image field 24 hours and 72 hours after seeding. Dots represent mean values and error bars represent standard deviation (2-way ANOVA with Tukey’s HSD; n = 3 per time point; each replicate is an average over 25 images). Scale bars = 50 µm. All images were acquired on the CX7 HCS platform with a 20x objective lens. For all graphs, * p< 0.05, ** p < 0.01, *** p < 0.001, **** < 0.0001. See also Figure S2 and Figure S3.

### H2B-fused fluorophores reveal increased cellular proliferation in PRKN knockout astrocytes

While dopaminergic neuron death is the predominant cellular outcome in Parkinson’s Disease (PD), other cell types in the brain may change cell cycle regulation under neurotoxic conditions. Tracking cell numbers to monitor proliferation, senescence, or death in vitro can help researchers understand how different cell populations are responding to neurotoxic stress or potential therapeutic strategies.

First, we validated that our nuclear-localized construct, H2B-Emerald, could be used to track cellular proliferation. The proliferation of H2B-Emerald transduced hiPSC-derived astrocytes was induced using fetal bovine serum (FBS) or blocked by inhibiting the proteasome with MG-132. Astrocyte proliferation was measured by changes in the number of H2B-Emerald nuclei. As expected, the astrocytes treated with FBS proliferated twice as rapidly as the untreated astrocytes over 48 hours (2-way ANOVA with Tukey’s HSD, p-adj = 0.005, **Fig. S2**), while the astrocytes treated with MG-132 did not significantly proliferate over the same period (adj-p = 0.08, **Fig. S2**). Thus, the GEM-SCOPe H2B-fluorophore fusion proteins can be used to track cellular proliferation over multiple days in live cell cultures.

To examine the effect of loss of *PRKN* on astrocyte proliferation, we transduced wild-type (*PRKN*^+/+^) astrocytes with H2B-Emerald and isogenic *PRKN* knockout (*PRKN*^-/-^) astrocytes (**Fig. S3A**) with H2B-mCherry (**Fig. 2D**). Leveraging the modularity of GEM-SCOPe, we were able to mix the two astrocyte populations together and track their proliferation simultaneously. Over 48 hours, the *PRKN*^-/-^ population increased significantly (2-way ANOVA with Tukey’s HSD, p-adj = 0.03; **Fig. 2E, F**) while the *PRKN*^+/+^ population did not proliferate in the same period (2-way ANOVA with Tukey’s HSD, p-adj = 0.4; **Fig. 2E, F**). Surprisingly, these results contradict mouse studies that found that PRKN-null mice had decreased astroglia proliferation^55^. However, PRKN has been widely described as a tumor suppressor, with loss-of-function mutations arising in a variety of cancers, including glioblastomas^56–58^. Studies on PRKN in the context of cancer have demonstrated that PRKN regulates the degradation of cyclins and cyclin-dependent kinases involved in the G1/S transition, the loss of which promotes unchecked proliferation^57,59,60^. It has previously been hypothesized that the outcome of loss-of-function PRKN mutations are highly cell-context dependent; the increased half-life of cyclins leads to cancer in mitotically competent cells while the same cyclins may promote apoptosis upon cell cycle reentry in post-mitotic cells, such as neurons^57,61–64^. Our results in human astrocytes highlight the importance of studying disease-relevant human cell types to capture the intricacies and nuances specific to the human brain that could contribute to PD.

### LAMP1-localized fluorophores reveal changes in lysosomal distribution in response to chemical and genetic stressors

Lysosomal dysfunction is widely implicated in PD and other forms of neurodegeneration. Several PD-risk and familial genes are involved in the autophagy-lysosomal-endosomal pathway^65–67^, and many hallmarks of lysosomal dysfunction associated with familial PD are present in idiopathic PD ^66,68,69^. Disruption to the autophagy-lysosomal-endosomal pathway is of broad importance in PD and there is a crucial need to study lysosomal function and dysfunction in the context of PD pathogenesis. As described above, we fused the N-terminus of LAMP1 to a fluorophore (Emerald or mCherry) to specifically target the fluorescent signal to the lysosome (**Fig. 1C, 3A**). To validate the GEM-SCOPe LAMP1-mCherry signal responds to lysosomal stress, hiPSC-derived astrocytes transduced with LAMP1-mCherry lentivirus were exposed to bafilomycinA1 (BafA1) for

15 hours. BafA1 inhibits lysosomal proton pump V-ATPases, which blocks autophagosome-lysosome fusion and lysosomal acidification, impairing lysosomal function ^70,71^. hiPSC-derived astrocytes were simultaneously transduced with a non-localized Emerald fluorophore lentivirus, to provide a cell body marker. We used CellProfiler ^72^ to evaluate the lysosomal number, size, and distribution within each cell. Compared to vehicle (DMSO) treatment, treatment with BafA1 increased the number of LAMP1-mCherry vesicles (unpaired t-test, p = 0.029; **Fig. 3B, C**), but did not affect lysosome size (unpaired t-test, p = 0.16, **Fig. 3B, C**). Treatment with BafA1 altered the distribution of lysosomes within the astrocytes. Astrocytes treated with BafA1 had a more dispersed distribution of LAMP1-mCherry signal, with only 40.0±2.0% concentrated in the perinuclear area, in contrast to 62.4±7.5% in vehicle-treated astrocytes (unpaired t-test, p = 0.0012; **Fig. 3B, C**). Perinuclear and juxtanuclear lysosomes have been posited to be more acidic than peripheral lysosomes^73^. Lysosomal distribution can also be affected by cytosolic pH, with acidification promoting the spreading of lysosomes away from the nucleus^74–76^. Since BafA1 inhibits V-ATPases responsible for establishing and maintaining lysosomal pH, the redistribution of lysosomes away from the nucleus upon BafA1 treatment might reflect a change in lysosomal or cytosolic pH, and therefore lysosomal function, in the treated astrocytes.

**Fig. 3:**
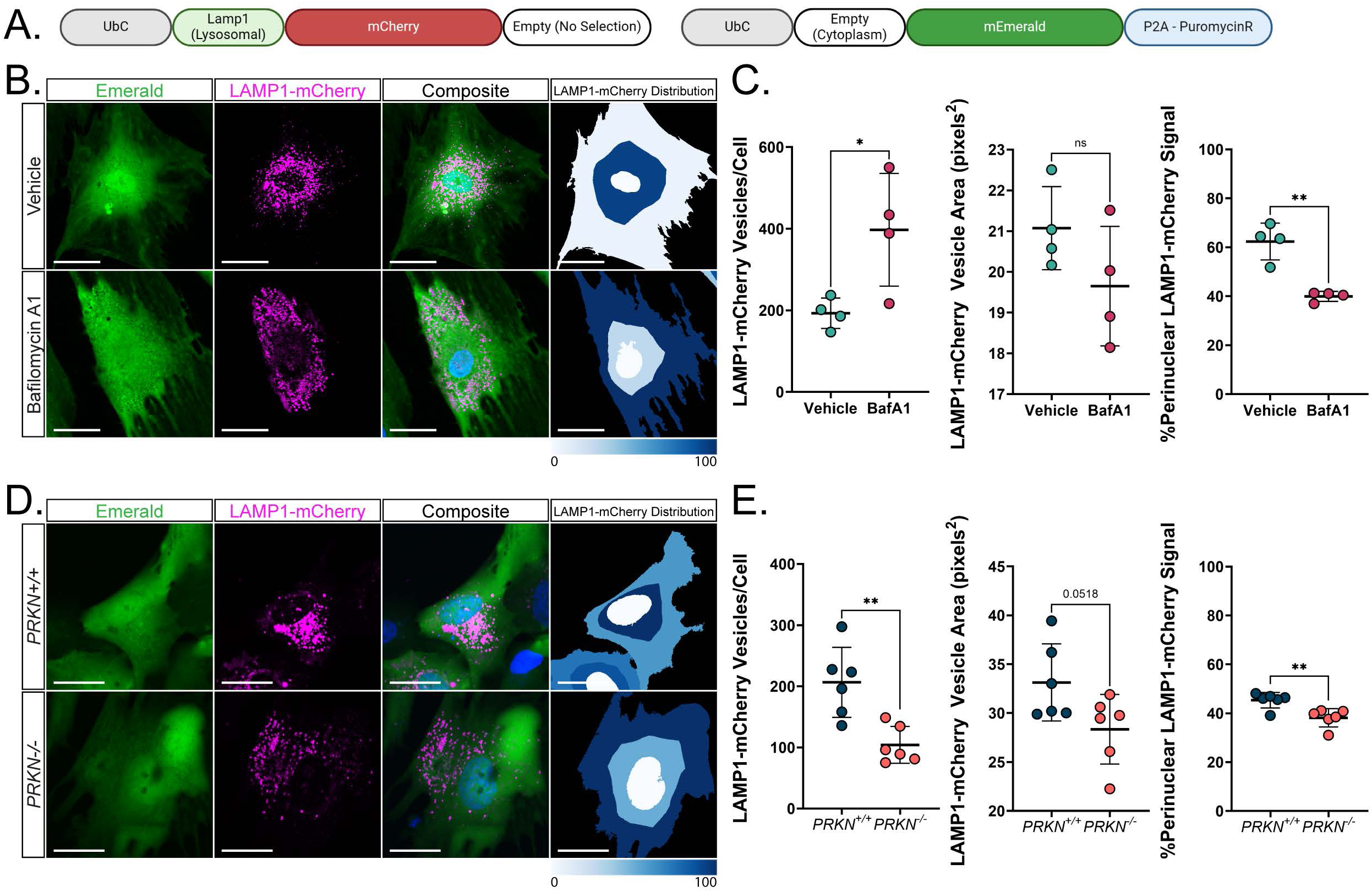
Lysosome-localized fluorophores reveal changes in lysosomal distribution upon chemical and genetic perturbation. **(A)** Schematic of the lentiviral constructs used in the following panels localizing (top) mCherry to the lysosome with the 5’ sequence of *LAMP1* and (bottom) Emerald to the whole cell by not including a localization sequence. **(B)** Representative images of astrocytes transduced with Emerald and LAMP1-mCherry lentiviruses and treated with 100 nM bafilomycin A1 (bottom) or a vehicle (DMSO; top). Nuclei were stained with Hoechst33342. Far right: Output from CellProfiler showing the distribution Lamp1-mCherry signal as a fraction of signal in each bin. Darker blue indicates a higher fraction of signal in that bin. Scale bars = 50 µm. **(C)** Quantification of lysosomal number per cell (left), average vesicle area (middle) and fraction of perinuclear lysosomes (right) using Lamp1-mCherry signal in vehicle (DMSO) and bafilomycin A1 (100 nM) treated astrocytes. Emerald was used to determine cell boundaries. Central bars represent mean and error bars represent standard deviation (unpaired t-tests; n = 4; each replicate is an average over 10 images). **(D)** Representative images of *PRKN^+/+^* (top) and *PRKN^-/-^* (bottom) astrocytes transduced with cytoplasmic Emerald and LAMP1-mCherry lentiviruses. Nuclei were stained with Hoechst33342. Far right: Output from CellProfiler showing the distribution Lamp1-mCherry signal as a fraction of signal in each bin. Darker blue indicates a higher fraction of signal in that bin. Scale bars = 25 µm. (**E**) Quantification of Lamp1-mCherry signal in *PRKN^+/+^* and *PRKN^-/-^* astrocytes to measure lysosomal number per cell (left), average vesicle area (middle) and fraction of perinuclear lysosomes (right). Emerald fluorescence was used to determine cell boundaries. Central bars represent mean and error bars represent standard deviation (unpaired t-tests; n = 6; each replicate is an average over 7 images). All images were acquired on a Nikon Ti2E AX R confocal microscope with a 20x objective lens and 6x optical zoom. For all graphs, * p< 0.05, ** p < 0.01, *** p < 0.001, **** < 0.0001.

In addition to its role in ubiquitinating damaged mitochondria for degradation via the autophagy-lysosomal pathway^31–33^, PRKN has been shown to regulate endosome organization^77^ and mitochondria-lysosome contact sites. Although it has been reported that *PRKN* knockout in hiPSC-derived dopaminergic neurons leads to an increase in lysosome number and size compared to isogenic, wild-type dopaminergic neurons^78^, the effect of *PRKN* knockout on lysosomes on human astrocytes is largely unknown. Due to PRKN’s role in regulating the autophagy-lysosomal pathway, we hypothesized *PRKN^-/-^* in human astrocytes would disrupt lysosomal dynamics. To test this hypothesis, we applied the same CellProfiler analysis on *PRKN*^+/+^ and *PRKN*^-/-^ hiPSC-derived astrocytes transduced with GEM-SCOPe Emerald and LAMP1-mCherry lentivirus. *PRKN*^-/-^ astrocytes had half as many LAMP1-mCherry vesicles per cell compared to *PRKN*^+/+^ astrocytes (unpaired t-test, p = 0.003, **Fig. 3D, E**). Additionally, the *PRKN*^-/-^ lysosomes were smaller than the *PRKN*^+/+^ lysosomes (unpaired t-test, p = 0.052, **Fig. 3D, E**). Finally, the *PRKN^-/-^* astrocytes had a decrease in perinuclear LAMP1-mCherry, with only 38±4% of LAMP1-mCherry signal located in the perinuclear area, compared to 45±3% in *PRKN*^+/+^ (unpaired t-test, p = 0.005, **Fig. 3D, E**). Our results utilizing our GEM-SCOPe LAMP1-mCherry fluorophore suggest that lysosomal number and distribution in astrocytes is altered upon loss of *PRKN*. However, further studies to examine lysosomal content and activity are needed to understand the mechanisms by which lysosomes are affected, the implications of this effect on neuron death, and how this could differ from observations in other cell types.

### Mitochondria-localized fluorophores capture mitochondrial fragmentation and alterations in mitochondrial trafficking along PRKN knockout dopaminergic neuron axons

Much like lysosomal dysfunction, mitochondrial dysfunction is implicated in PD and neurodegeneration. Therefore, monitoring mitochondrial network dynamics in response to stimuli and stress is important in understanding neurodegenerative diseases. As described above, we fused the N-terminus of the mitochondrial protein COX8A to a fluorophore (Emerald or mCherry) to localize the fluorophore specifically to the mitochondria (**Fig. 1C, 4A**). To validate that GEM-SCOPe COX8A localized fluorophores reflect changes to mitochondrial morphology, we induced mitochondrial stress by treating hiPSC-derived astrocytes transduced with the COX8A-mEmerald lentivirus with 2µM oligomycin for 4 hours. Mitochondrial stress induces fragmentation of the mitochondrial network, which facilitates mitophagy, but also decreases mitochondrial network respiratory capacity^79,80^. Oligomycin, an inhibitor of ATP synthase, is a known inducer of mitochondrial fragmentation. Treatment with oligomycin induced mitochondrial fragmentation when compared to vehicle-treated astrocytes. The average area of mitochondria was reduced from 1.04 µm^2^ in vehicle-treated astrocytes to 0.80 µm^2^ in oligomycin-treated astrocytes (1-way ANOVA with Dunett’s Test, p-adj = 0.0002; **Fig. 4B, C left)**, and average branch length was reduced from 1.28 µm in vehicle-treated astrocytes to 0.90 µm in oligomycin treated astrocytes (1-way ANOVA with Dunett’s Test, p-adj < 0.0001; **Fig. 4B, C middle)**, as expected from a more fragmented mitochondrial network. The aspect ratio is the ratio of the mitochondrial length to width. An aspect ratio of 1 indicates a perfect circle, while larger aspect ratios indicate longer and narrower shapes. The mean aspect ratio decreased from 2.52 in vehicle-treated astrocytes to 2.14 in oligomycin-treated astrocytes (1-way ANOVA with Dunett’s Test, p-adj < 0.0001; **Fig. 4B, C right**), another indicator that mitochondria are fragmenting upon oligomycin treatment. Metrics of mitochondrial network fragmentation (mitochondrial area, branch length, and aspect ratio) are all affected in a dose-dependent manner, with increasing concentrations of oligomycin having increasingly more severe effects on mitochondrial network fragmentation (**Fig. S4**).

**Fig. 4:**
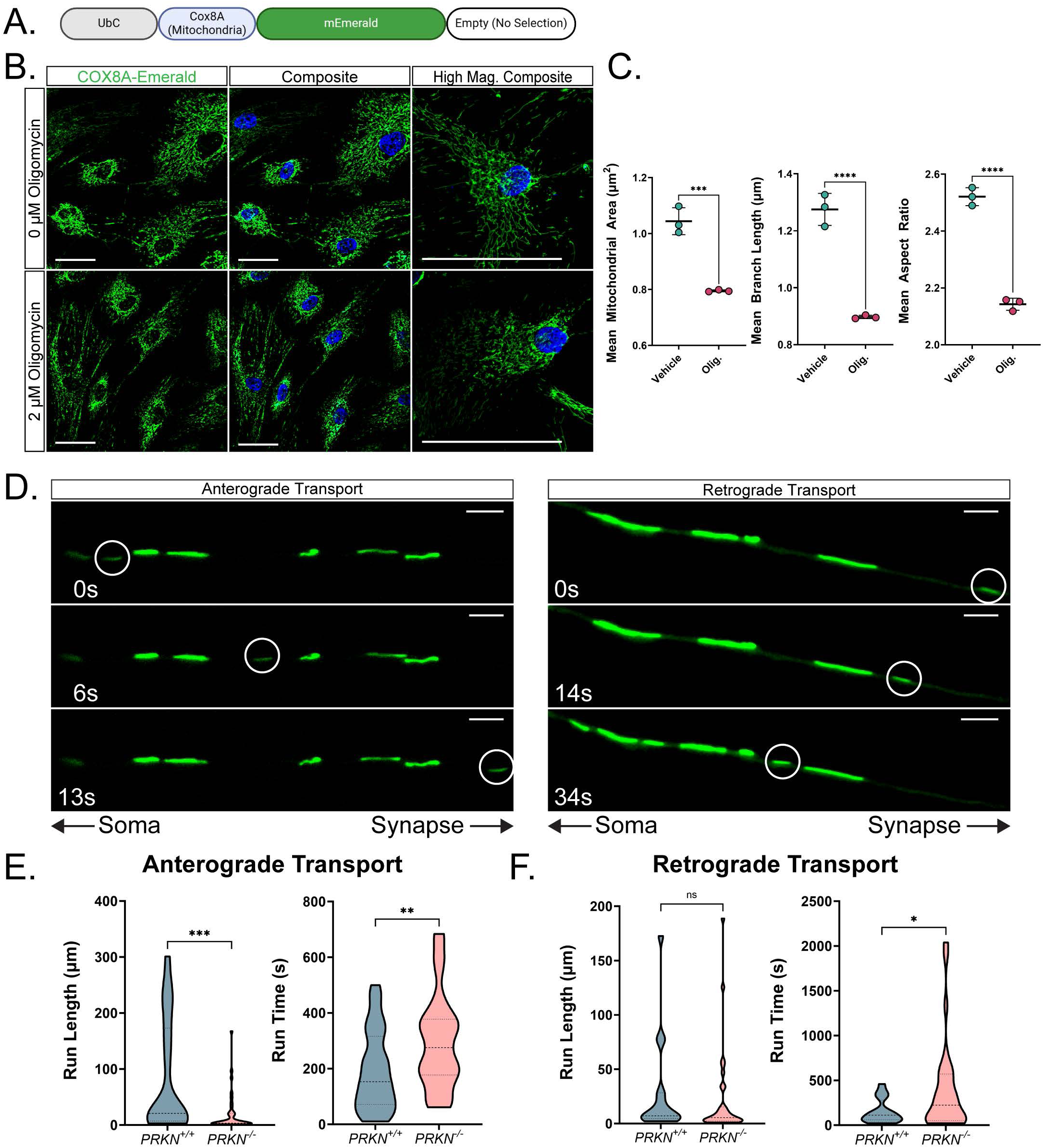
Mitochondria-localized fluorophores resolve changes in mitochondrial network dynamics. **(A)** Schematic of the lentiviral construct used in the following panels localizing Emerald to the mitochondria with the targeting sequence of COX8A. **(B)** Representative images of astrocytes transduced with COX8A-Emerald and treated with a vehicle (DMSO; top) or 2 uM oligomycin (bottom). Nuclei were stained with Hoechst33342. High magnification images (far right) highlight the changes to the mitochondrial network upon oligomycin treatment. Scale bars = 50 µm. **(C)** Quantification of mean mitochondrial area, mean mitochondrial branch length, and mean aspect ratio upon oligomycin treatment. Central bars represent mean and error bars represent standard deviation (1-way ANOVAs with Dunett’s test; n = 3; each replicate is an average over 36 images) **(D)** Representative images of mitochondria labeled with COX8A-Emerald moving along *PRKN^-/-^* neuron axons in the anterograde (left) and retrograde (right) directions. Scale bars = 5 µm. Images were taken on a Leica DMi8 microscope for 1 hour per field **(E-F**). Quantification of mitochondrial run length (left) and run time (right) of COX8A-Emerald labeled mitochondria in *PRKN^+/+^* and *PRKN^-/-^* dopaminergic neuron axons moving anterograde (E) and retrograde (F) as determined by kymograph analysis (unpaired t-test or Welch’s test; average of n = 37.5 organoids for anterograde and 23.5 organoids for retrograde; each replicate is an average over 1-3 fields). See also Figure S4. For all graphs, * p< 0.05, ** p < 0.01, *** p < 0.001, **** < 0.0001.

The transport of mitochondria along neuronal axons is critical for proper synaptic function and neurotransmitter release. We hypothesized *PRKN*^-/-^ may impair axonal mitochondrial transport. To investigate this hypothesis, we transduced wild-type (*PRKN^+/+^*) and *PRKN* knock-out (*PRKN^-/-^*) hiPSC-derived midbrain organoids with COX8A-mEmerald. To facilitate imaging of axonal mitochondrial transport, we generated radial axonal arbors by splatting organoids onto the surface of a plate and allowing the axons to grow out^81^. On day 164 of organoid differentiation, fluorescent mitochondria were imaged at high magnification over several minutes to monitor the movement of mitochondria along these axons (**Fig. 4D; Video S1**). In the anterograde direction (moving away from the soma), *PRKN^-/-^* mitochondria moved only 20% of the distance covered by *PRKN^+/+^* mitochondria (Welch’s t-test, p < 0.0001, **Fig. 4E left**) and took 50% longer (unpaired t-test, p = 0.005; **Fig. 4E right**). Retrograde-moving mitochondria moved the same distance (unpaired t-test, p = 0.59, **Fig. 4F left)** but the *PRKN^-/-^* mitochondria moved for 150% longer compared to the *PRKN^+/+^* mitochondria (Welch’s t-test, p = 0.02; **Fig. 4F right**). PRKN has previously been implicated in mitochondrial transport due to its ubiquitination of mitochondrial motor adaptor complexes^82^. Our results demonstrate that PRKN loss of function impairs mitochondrial trafficking in human neurons, likely reducing the delivery of functional mitochondria synaptic extremities and ultimately compromising neuronal function.

### Dynamic fluorophores measure deficiencies in mitochondrial turnover and glutathione reduction potential in PRKN-knockout astrocytes

We demonstrated the utility of single excitation and emission wavelengths to discern phenotypes regarding cellular and subcellular dynamics. Numerous groups have developed fluorophores that change emission wavelengths under certain conditions or circumstances. We introduced several multi-emission fluorophores into GEM-SCOPe to enable quantification of organelle turnover, acidification, and oxidation, in addition to the localization information described with the previous fluorophores.

Fluorescent Timer is a mutated dsRed protein that irreversibly shifts its fluorescence emission from green to red due to tyrosine oxidation and can be used as a molecular clock to track protein turnover (**Fig. 5A**).^22^ We, therefore, localized GEM-SCOPe-Timer to the mitochondria (COX8A-Timer), to further investigate mitochondrial turnover^83,84^ (**Fig. 5B**). To first validate COX8A-Timer, mitophagy was inhibited by treating hiPSC-derived astrocytes transduced with COX8A-Timer with the V-ATPase inhibitor bafilomycinA1 (BafA1). BafA1 treatment led to a significant increase in the ratio of red-fluorescent mitochondria over green-fluorescent mitochondria compared to vehicle control astrocytes (unpaired t-test, p < 0.0001; **Fig. 5C, D**). Since mitochondrial degradation and turnover occur primarily through mitophagy, inhibition of mitophagy and mitochondrial recycling via BafA1 treatment results in a predicted increase in longer-lived mitochondria.

Because PRKN is directly involved in mitophagy, and we observed impaired retrograde mitochondrial transport in *PRKN^-/-^* neurons, we wanted to measure changes in mitochondrial turnover in human *PRKN^-/-^* cells. We transduced wild-type (*PRKN^+/+^*) and *PRKN* knockout *(PRKN^-/-^*) hiPSC-derived astrocytes with GEM-SCOPe COX8A-Timer. The ratio of red-fluorescent signal over green-fluorescent signal significantly increased in *PRKN^-/-^*astrocytes compared to *PRKN^+/+^*astrocytes (unpaired t-test, p = 0.026, **Fig. 5E, F**). When broken down by individual channel, there was no change in the green fluorescence (2-way ANOVA with Sidak correction, p-adj = 0.1288, **Fig. 5G**), indicating no change in mitochondrial biogenesis. The increase in red to green ratio was driven by a significant increase in red-fluorescent signal (2-way ANOVA with Sidak correction, p-adj = 0.0005, **Fig. 5G**), which is in concordance with an accumulation of older mitochondria due to a failure to properly degrade mitochondria in the absence of PRKN. This observation is supported in a Parkin-knockout mouse that showed a subtle increase in the protein half-life, and therefore reduced protein turnover, of the subunits of the electron transport chain compared to wild-type mice^85^.

An expected consequence of impaired mitochondrial recycling is an accumulation of damaged mitochondria and an increase in mitochondria-associated reactive oxygen species (ROS) from inefficient cellular respiration. To measure reactive oxygen levels in live astrocytes, we introduced GRX1-roGFP2, an established fluorophore for measuring ROS, into GEM-SCOPe^25^. GRX1-roGFP2 is a fusion protein between GRX1 and roGFP2. When GRX1 is oxidized by glutathione, it can subsequently oxidize roGFP2. Oxidized roGFP2 shifts its emission spectrum from green fluorescence to blue fluorescence (**Fig. 6A**)^25,86^. Thus, the ratio of green to blue fluorescence is an indicator of glutathione oxidation state. When GRX1-roGFP2 is further coupled with a mitochondrial localization sequence, we can measure the oxidative stress of the mitochondria.

**Fig. 5:**
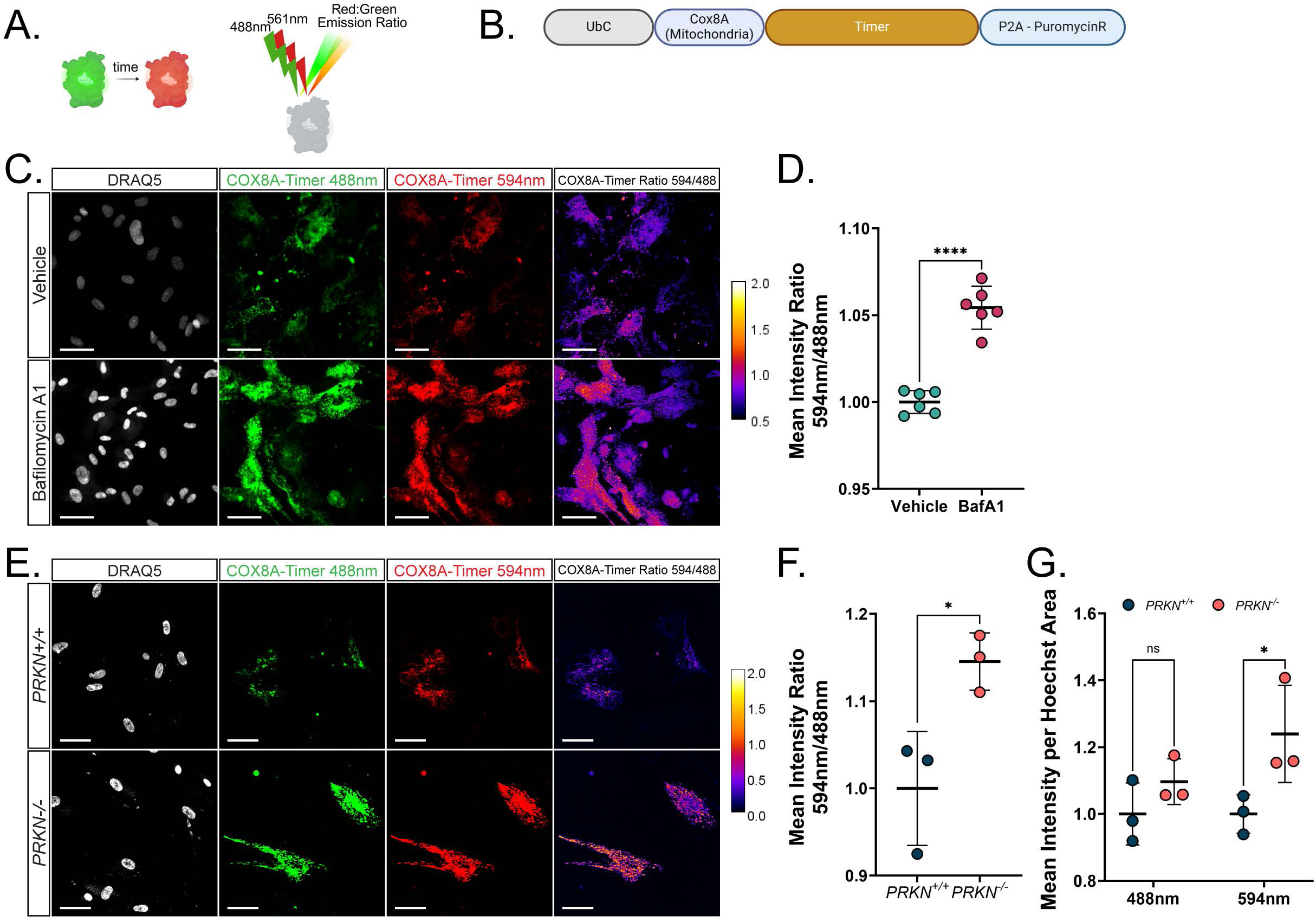
Timer fluorophore measures changes in subcellular mitochondrial turnover in *PRKN^-/-^*(A) Schematic demonstrating how the fluorescent protein Timer works. Timer undergoes a fluorescent shift from a green emission to a red emission overtime; analysis of Timer relies on examining the ratio between the red and green emission. **(B)** Schematic of the lentiviral construct used in the following panels localizing Timer to the mitochondria by using the targeting sequence of COX8A. **(C)** Representative images of astrocytes transduced with COX8A-Timer and treated with a vehicle (DMSO; top) or 100 nM bafilomycin A1 (bottom) for 24 hours. Images were acquired with a 488-excitation line and a 594-excitation line. Far right: ratiometric representation of the red channel divided by the green channel and pseudo colored so that orange, yellow and white indicate more relative red emission and black and purple indicate more relative green emission. Nuclei were stained with DRAQ5. Scale bars = 50 µm. **(D)** Quantification of the ratio of mean intensity of emission from excitation with a 594nm laser to the mean intensity of emission from excitation with a 488 nm laser in astrocytes expressing COX8A-Timer and treated with a vehicle (DMSO) or 100 nM bafilomycin A1 for 24 hours. Central bars represent mean and error bars represent standard deviation (unpaired t-test; n = 6; each replicate is an average over 25 images). (**E**) Representative images of *PRKN^+/+^* and *PRKN^-/-^* astrocytes transduced with COX8A-Timer. Images were acquired with a 488-excitation line and a 594-excitation line. Far right: ratiometric representation of the red channel divided by the green channel and pseudo colored so that orange, yellow and white indicate more relative red emission and black and purple indicate more relative green emission. Nuclei were stained with DRAQ5. Scale bars = 50 µm. **(F)** Quantification of the ratio of mean intensity of emission from excitation with a 594nm laser to the mean intensity of emission from excitation with a 488 nm laser in *PRKN^+/+^* and *PRKN^-/-^* astrocytes expressing COX8A-Timer. Central bars represent mean and error bars represent standard deviation (n = 3; each replicate is an average over 25 images). **(G)** Quantification of mean intensity normalized by nuclear area of green and red fluorescence in *PRKN^+/+^* and *PRKN^-/-^*astrocytes expressing COX8A-Timer. Central bars represent mean and error bars represent standard deviation (2-way ANOVA with Sidak correction; n = 3; each replicate is an average over 25 images). All images were acquired on a CX7 HCS platform with a 20x objective lens. For all graphs, * p< 0.05, ** p < 0.01, *** p < 0.001, **** < 0.0001.

**Fig. 6:**
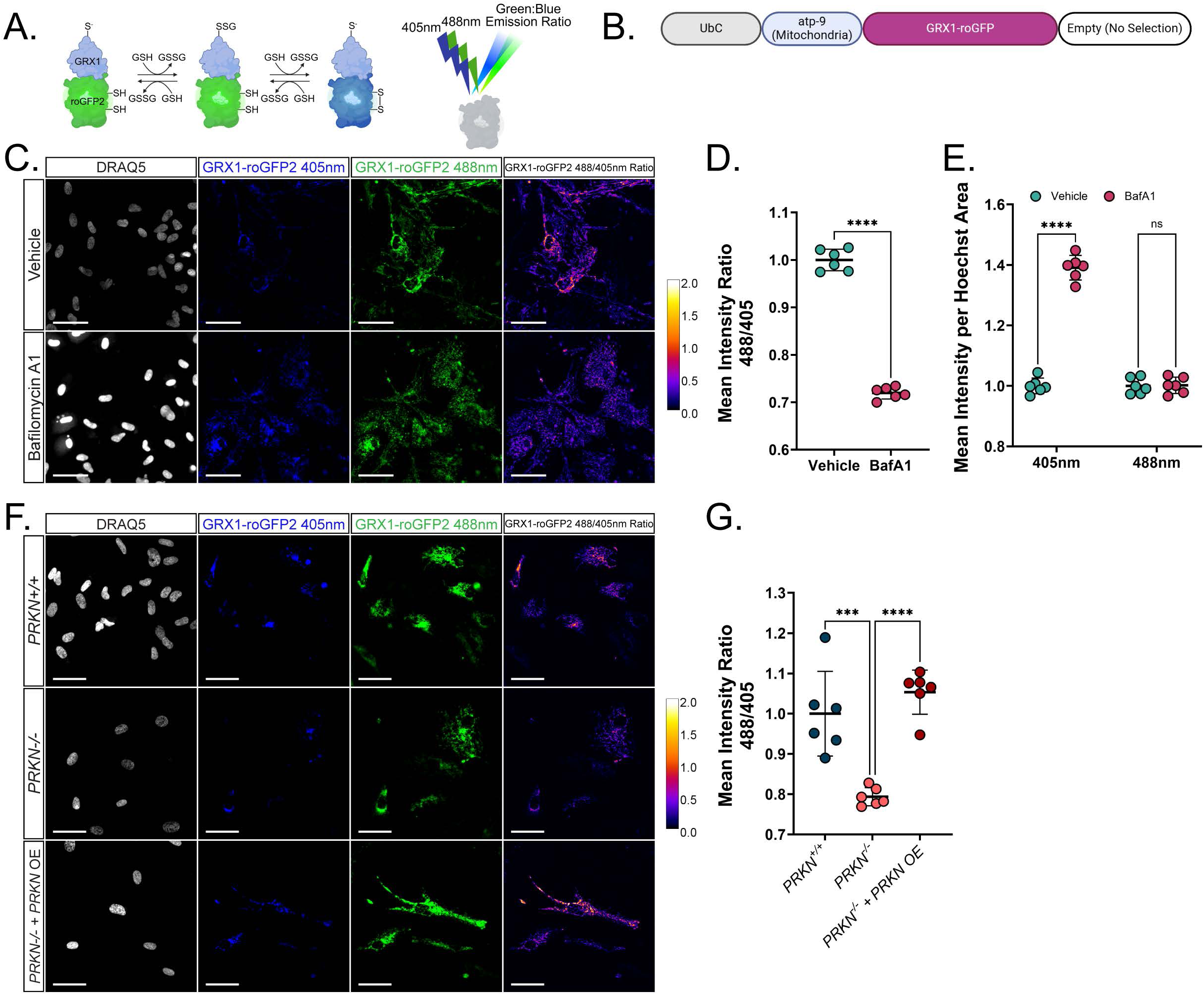
GRX1-roGFP2 fluorophore is used to quantify glutathione oxidation in chemical and genetic models of oxidative stress. **(A)** Schematic demonstrating how the fluorescent protein fusion Grx1-roGFP2 works. Oxidation of Grx1 upon the oxidation of glutathione results in the eventual oxidation of roGFP2, shifting its emission spectrum from green to blue. **(B)** Schematic of the lentiviral construct used in the following panels localizing Grx1-roGFP2 to the mitochondria. This fluorophore was localized using atp-9 instead of COX8A. **(C)** Representative images of astrocytes transduced with mito-GRX1-roGFP2 and treated with a vehicle (DMSO; top) or 100 nM bafilomycin A1 (bottom) for 24 hours. Images were acquired with a 405-excitation laser and a 488-excitation laser. Far right: ratiometric representation of the green channel divided by the blue channel and then pseudo colored so that orange, yellow and white indicate more relative green emission while black and purple represent more blue emission. Nuclei were stained with DRAQ5. Scale bars = 50 µm. **(D)** Quantification of the ratio of mean intensity from 488-excitation to the mean intensity from 405 excitation. In astrocytes expression mito-Grx1-roGFP2 and treated with a vehicle or 100 nM bafilomycin A1. Central bars represent mean and error bars represent standard deviation (unpaired t-test; n = 6; each replicate is an average over 25 images). **(E)** Quantification of mean intensity normalized by nuclear area of blue and green fluorescence in vehicle and bafilomycin A1 treated astrocytes expressing mito-Grx1-roGFP2. Central bars represent mean and error bars represent standard deviation (n = 6; each replicate is an average of 25 images). **(F)** Representative images of *PRKN^+/+^*, *PRKN^-/-^*, and *PRKN^-/-^* with *PRKN*-overexpression astrocytes transduced with mito-Grx1-roGFP2. Images were acquired with a 405-excitation laser and a 588-excitation laser. Far right: ratiometric representation of the green channel divided by the blue channel and pseudo colored so that orange, yellow and white indicate more relative green emission and black and purple indicate more relative blue emission. Nuclei were stained with DRAQ5. Scale bars = 50 µm. **(G)** Quantification of the ratio of mean intensity of emission from excitation with a 488nm laser to the mean intensity of emission from excitation with a 405 nm laser in *PRKN^+/+^*, *PRKN^-/-^*, and *PRKN^-/-^* with *PRKN*-overexpression astrocytes expressing mito-Grx1-roGFP2. Central bars represent mean and error bars represent standard deviation (1-way ANOVA with Tukey’s HSD; n = 6; each replicate is an average over 36 images). For all graphs, * p< 0.05, ** p < 0.01, *** p < 0.001, **** < 0.0001. See also Figure S3.

This construct was available as a retroviral vector using the signal sequence from *Neurospora crassa* ATP synthase protein 9^25^. When we cloned it into the FUW lentivirus backbone, we decided to keep that localization sequence rather than replace it with the signal sequence for COX8A used in the other GEM-SCOPe mitochondria-localized fluorophores (**Fig. 6B**). To validate GEM-SCOPe GRX1-roGFP2 was working as expected, hiPSC-derived astrocytes transduced with mito-GRX1-roGFP2 were treated with BafA1 to induce oxidative stress^87^. Astrocytes treated with BafA1 had a significant decrease in the ratio of green fluorescence to blue fluorescence when compared to vehicle (DMSO) treated astrocytes (unpaired t-test, p < 0.0001, **Fig. 6C, D**). The effect is driven by a significant increase specifically in blue fluorescence (2-way ANOVA with Sidak correction, p-adj < 0.0001, **Fig. 6E**) while the green fluorescence remained unchanged (2-way ANOVA with Sidak correction, p-adj = 0.99, **Fig. 6E**). The increase in blue fluorescence is due to an increase in roGFP2 oxidation, which is indicative of an increase in glutathione oxidation and the presence of reactive oxygen species. Thus, GEM-SCOPe-mito-GRX1-roGFP can be used to monitor glutathione reduction potential and how cells respond to reactive oxygen stress.

Because astrocytes play important metabolically supportive roles in the CNS and we observed PRKN*-*related impairments to mitochondrial recycling with COX8A-Timer (**Fig. 5E-G**), we wanted to understand if PRKN deficiency has effects on ROS production in astrocytes. We transduced *PRKN^+/+^*and *PRKN^-/-^* astrocytes with mito-GRX1-roGFP2 lentivirus. Without introducing any additional chemical mitochondrial stressors, we observed a significant reduction in the ratio of green-fluorescence signal to blue-fluorescence signal in *PRKN^-/-^* compared to *PRKN^+/+^* astrocytes (1-way ANOVA with Tukey’s HSD, p = 0.0003; **Fig. 6F, G**). This is indicative of an increase in ROS in *PRKN^-/-^*astrocytes which correlates with our other observation of decreased mitochondrial turnover in the same cells. We reintroduced the *PRKN* coding sequence into *PRKN*^-/-^ astrocytes via lentivirus (**Fig. S3B**), and found the ratio of green to blue fluorescence increased to levels comparable to *PRKN^+/+^* astrocytes (1-way ANOVA with Tukey’s HSD, p = 0.401); **Fig. 6F, G**), confirming that the increase in ROS observed in *PRKN^-/-^* astrocytes was due to the loss of *PRKN*. It has been previously demonstrated that *PRKN^-/-^* hiPSC-derived dopaminergic neurons show increased cellular ROS^15^ and *PRKN^-/-^* mice have decreased antioxidant capabilities and increased ROS-related protein and lipid damage^88^. By introducing fluorophore-biosensors into our lentiviral catalog, we were able to expand the phenotypes we can observe in live cells to measure changes in the intracellular environment in response to chemical and genetic perturbations.

## Discussion

Here, we developed and validated GEM-SCOPe (Genetically Engineered, Modular, SubCellular Organelle Probes), a collection of subcellularly targeted fluorophores encoded in a lentiviral backbone. By exploiting the modular design of the constructs, we easily cloned fluorophores with different excitation/emission spectra specified to localize to the nucleus, lysosomes, or mitochondria. We demonstrated that these fluorophores specifically localize to the desired organelle and can quantitatively assess cellular responses to chemical perturbations. Finally, we applied GEM-SCOPe to human *PRKN*-knockout astrocytes and neurons, which revealed widespread perturbations to cellular proliferation, lysosomal distribution, and mitochondrial dynamics, providing new insights into PD pathogenesis.

Gliosis has been widely reported in human post-mortem brain studies of patients with *PRKN* loss-of-function mutations^89–92^. Similar pro-inflammatory phenotypes have also been observed in mouse models and stem-cell-derived glial models of PRKN deficiency^93,94^. However, there is still a limited understanding of how genetic mutations related to PD modify astrocytes and their role in neurodegeneration. We used GEM-SCOPe to investigate cellular, mitochondrial, and lysosomal dynamics in astrocytes lacking functional PRKN protein. Using nucleus-localized fluorophores, we show that *PRKN* knockout human astrocytes have increased cellular proliferation compared to isogenic, wild-type astrocytes. LAMP1-localized fluorophores revealed that lysosomes in *PRKN* knockout astrocytes are distributed further from the perinuclear nuclear area. Finally, a variety of mitochondria-localized fluorophores revealed that *PRKN* knockout astrocytes have impaired mitochondrial transport, decreased mitochondrial turnover, increased reactive oxygen species, and oxidized glutathione. One limitation to the *PRKN* knockout model is the genetic variability that can arise from culturing cells in vitro. While additional experiments must be done to support these findings, our goal was to demonstrate the utility and efficacy of GEM-SCOPe to detect measurable genetically mediated differences in cell models. GEM-SCOPe highlights that there is extensive dysregulation and disruption to normal organelle dynamics in human *PRKN* astrocytes that warrants further investigation.

GEM-SCOPe was developed in a 3^rd^ generation lentiviral system. We selected this delivery system because 1) lentiviruses are safe to produce; 2) they infect both dividing and non-dividing cells; and 3) they can be used to develop stable cell lines. 3^rd^ generation lentiviral systems are replication-incompetent, so cells that are transduced with lentivirus cannot produce more viral particles. Only the transfer plasmid, which contains the insert of interest, can be integrated into the host genome; the other plasmids that encode lentiviral structural and envelope genes do not integrate into the host genome, preventing viral replication. Most lentiviruses are produced with VSV-G (Vesicular stomatitis virus G glycoprotein) as the envelope protein, as it provides broad tropism and enables high transduction efficiency in a wide variety of cell types^95^. Finally, GEM-SCOPe lentiviruses were developed with the option to include an antibiotic-resistance gene (**Fig. 1A**), which can be used to establish stable cell lines that have incorporated the lentiviral load.

Although lentiviruses provide a lot of flexibility and safety when delivering genetic information, they also possess some limitations. Lentiviruses have an ideal plasmid size of 9-10kb, and viral titer can decrease if plasmids exceed this size^96^. The H2B-mTagBFP2, H2B-mCherry, and H2B-Emerald lentiviruses (**Fig. 1, 2**) are about 11kb each and transduced hiPSC-derived astrocytes with about 90% efficiency. Coupled with the puromycin resistance that was also included in this construct, we were still able to generate a population with 100% viral transduction despite the size limitation. Another drawback of lentiviral gene delivery is that inserts often integrate into actively transcribed loci^97^. Differentiation of hiPSCs into specific cell types requires extensive chromatin remodeling, and genomic sequences integrated via lentivirus can often be silenced in this process. In the case where hiPSCs must be transduced before differentiation, antibiotic selection during differentiation can select for cells that have integrated the viral load into a genomic region that is not silenced. The final drawback of lentiviral gene delivery is that fluorescence intensity is dependent on the viral copy number in each cell and cannot be used as a readout for organelle abundance. Rather, we need to use measurements like distribution, size, and ratios of different emissions within the same cell. This is not just a limitation of lentiviral gene delivery but of other viral and non-viral delivery systems.

In this study, we focused on PD and specifically the alterations to astrocytes with *PRKN* mutations. However, alterations in proliferation, mitochondria, and lysosomes are by no means unique to *PRKN* or PD. Many reviews cover the extensive roles of mitochondria and lysosomes in health and disease^98–101^. GEM-SCOPe can be applied to countless cell types and *in vitro* disease models to investigate subcellular dynamics in the development and progression of a variety of diseases, beyond neurodegeneration. Due to its modular design, GEM-SCOPe can be expanded and molded by research groups to meet specific needs. Different targeting sequences can be integrated to localize fluorophores to relevant subcellular compartments. New fluorophores and biosensors are always being developed to measure additional intracellular conditions. By applying GEM-SCOPe to disease models, we will be able to better understand how cells are disrupted over the course of disease progression, providing insight for future therapeutic targets.

## Resource Availability

### Lead contact

Further information and requests for resources and reagents should be directed to and will be fulfilled by the lead contact, Joel W. Blanchard.

### Materials availability

All plasmids generated in this study have been deposited to Addgene (**Table S1**).

### Data and code availability

- Raw data from all Figures and Supplemental Figures is available on Mendeley (DOI: <u>^1^0.17632/fmxdjgrjw5.^1^</u>)
- This paper does not report original code.
- Any additional information required to reanalyze the data reported in this paper is available from the lead contact upon request.

### Limitations of Study

In this study, we focused on *PRKN^-/-^* astrocyte phenotypes grown in monocultures. I future studies, it would be crucial to examine astrocyte phenotypes in a more complex multi-culture system, as responses to inputs from other cell types may affect astrocyte behavior in a disease-relevant manner. Furthermore, we only used a *PRKN^-/-^* cell line generated from one human, male donor. In future studies, it would be important to validate our findings in other iPSC lines, especially from both male and female donors, to confirm the relevance of observed phenotypes across cell lines with different genetic backgrounds.

## Acknowledgements

The authors thank Deanna Benson for discussions on mitochondrial transport and kymograph analysis. We thank the Microscopy CoRE at Icahn School of Medicine for providing access to the Leica DMi8 and Thermo Scientific CX7 High Content Screening Platform, as well as training and technical expertise for using these microscopes.

This work has been funded in part with federal funds from NASA under contract #80ARC022CA004 titled “Identification of Biomarkers and Pathological Mechanisms via Longitudinal Analysis of Neurological and Cerebrovascular Responses to Neurotoxic Stress Using a Multi-cellular Integrated Model of the Human Brain.” This research was funded in whole or in part by Aligning Science Across Parkinson’s (ASAP-024297) through the Michael J. Fox Foundation for Parkinson’s Research (MJFF). For open access, the authors have applied a CC BY public copyright license to all Author Accepted Manuscripts arising from this submission. This work was also supported by funding from the NIH/NINDS (R01NS114239 and UH3NS115064), and the CureAlz Fund. C.G. was supported by funding from the NIH/NIA (T32AG04968) and NIH/NINDS (F31NS13090), and L.S. was supported by the Training Program in Stem Cell Biology fellowship from the New York State Department of Health (NYSTEM-C32561GG).

## Author Contributions

Conceptualization, C.G., T.A., and J.W.B.; Methodology, C.G., L.S., and T.A.; Validation, C.G.; Formal Analysis, C.G.; Investigation, C.G., T.K., L.S., and A.S.; Resources, C.G., T.K., L.S., and B.R.S.; Writing – Original Draft, C.G. and J.W.B.; Writing – Review & Editing, C.G., L.S., B.R.S., and J.W.B.; Visualization, C.G.; Funding Acquisition, T.A. and J.W.B.

## STAR Methods

### Experimental model and study participant details

The human cell lines used in this study are detailed in the key resources table as well as below. HEK293FT (RRID: CVCL_6911, fetal kidney origin, female sex) were maintained in DMEM (Gibco 10566) supplemented with 10% bovine calf serum (Cytiva SH30073) at 37 °C in a 5% CO2 incubator. Cells were passaged every 4-5 days, when they reached 90% confluency.

H1(WA01) hESCs (male), BJ-SiPS-D iPSCs (male), and AG09173 iPSCs (female) were maintained in StemFlex medium (Gibco A3349401) at 37 °C in a 5% CO2 incubator. Cells were passaged every 3-4 days, when they reached 90% confluency. WA01 *PRKN^-/-^* isogenic lines were previously generated and validated in Ahfeldt et. al, 2020.^15^ WA01 astrocytes were used for all PRKN-genotype dependent experiments. BJ-SiPS-D and AG09173 were used for organelle localization validation and chemical perturbation validation. All iPSC lines have undergone quality control analysis including tests for pluripotency, genomic instability, sterility, and cell line identity, and routine mycoplasma testing prior to experimentation.

### Method Details

Detailed methods are available at dx.doi.org/10.17504/protocols.io.4r3l2pj4pg1y/v1

### Molecular cloning

All vectors were derived from pFUW (Addgene plasmid #14882). pFUW was linearized using NheI-HF (NEB R3131) and BamHI-HF (NEB R3136) per manufacturer’s protocol, and the desired fragment was gel-purified (Macherey-Nagel 740609). Components of the insert were PCR amplified using Q5-Hot Start DNA Polymerase (NEB M0493) per manufacturer’s recommendations. Fluorophores were amplified from commercially available plasmids (see Resource Table for complete list) and localization sequences were amplified from cDNA. Primers were designed using the NEBuilder Assembly Tool (nebuilder.neb.com) to add 5’ and 3’ sequences that make the fragments compatible for Gibson assembly and ordered from IDT (Integrated DNA Technologies). Gibson reactions were set up with NEBuilder HiFi DNA Assembly Master Mix (NEB E2621), 50ng of linearized backbone, and 3-fold molar excess of each PCR fragment. For fragments less than 200bp, 5-fold molar excess was used. Reactions were incubated at 50°C for 60 minutes. The Gibson product was transformed in 10-beta competent *E. coli* (NEB C3019) grown on LB Agar plates with 100ug/mL ampicillin. Plasmids from single clones were isolated via miniprep (Macherey-Nagel 740499) and sequenced to confirm successful cloning.

### Lentivirus Production

Lentivirus was produced by transfection of HEK293T cells as adapted from Dull et al. 1998 ^102^. HEK293T cells were grown in Dulbecco’s Modified Eagle Medium (DMEM; Gibco 11965092) with 10% bovine calf serum (Cytiva SH30073) at 37 °C in a 5% CO_2_ incubator. The day before transfection, ∼10^7^ cells were plated per 15cm dish coated in 0.1% gelatin. Media was changed just prior to transfection. For one 15cm dish, 22.5ug of transfer vector was combined with 14.7ug of pMDLg/pRRE (Addgene plasmid #12251), 5.7ug of pRSV.Rev (Addgene plasmid #12253), and 7.9ug of pMD2.G (Addgene plasmid #12259) in a 15mL conical tube. The plasmid mixture was added to 1mL of 278 uM CaCl_2_ (Sigma C1016) and mixed thoroughly before adding 1mL of 2x BBS solution [280 mM NaCl (Fisher Scientific S271-1), 50 mM BES (Millipore 391334), 1.5mM Na_2_HPO_4_ (Sigma S5136); pH = 6.95] dropwise. Tubes were mixed by inversion, incubated at room temperature for 1 min, and then added dropwise to HEK293T dish. Cell culture media was changed after 16-24 h. Conditioned media was then collected every 24 h for 2 days; conditioned media from the same plate was combined over the two collections. Conditioned media was centrifuged at 2000 x g for 10 minutes to pellet debris and filtered through 0.22-um-pore-size PES filter (Millipore SCGP00525). To concentrate virus, filtered media was incubated with PEG 8000 (Sigma P2139) and NaCl, at a final concentration of 5% and 0.15M respectively, overnight at 4°C, then centrifuged at 3000 x g for 20 min. The pellet was resuspended in DMEM/F12 with GlutaMAX (Gibco 10565018) to 1% of the original supernatant volume. The concentrated virus was stored in aliquots at −80°C.

### Midbrain Organoid and Astrocyte Differentiation

hiPSC-derived astrocytes were extracted from 100-day-old midbrain organoids as described in Sarrafha et al 2021 and Parfitt et al 2024 ^81,103^. A detailed protocol can be found at https://doi.org/10.1016/j.xpro.2021.100463. In short, hiPSCs were cultured in StemFlex medium (Gibco A3349401) at 37 °C in a 5% CO_2_ incubator. hiPSCs were seeded into 125-mL spinner flasks (Corning 3152) and allowed to self-aggregate in StemFlex (Gibco A3349401) supplemented with 10 µM Y-27632 (Tocris 1254) and 1% Penicillin-Streptomycin (Gibco 15140122). Once aggregates were between 300 µm and 500 µm, differentiation was initiated by dual-SMAD inhibition with 10 µM SB431542 (Stemgent 04-0010), 100nM LDN193189 (Tocris 6053), 1x B27 without Vitamin A (Gibco 12587010) and 1x N2 (Gibco 17502048) in DMEM-F12 with GlutaMAX (Gibco 10565018). Midbrain patterning was achieved with the addition of 3 µM CHIR99021 (Tocris 4423), 2 µM purmorphamine (Stemcell 72204), and 1 µM SAG (Cayman 11914) starting 4 days after neural induction. After patterning, media was changed to DMEM/F12 with GlutaMAX, supplemented with 1x N2, 1x B27 without Vitamin A, 20 ng/mL GDNF (Peprotech 450-10), 20 ng/mL BDNF (Peprotech 450-02), 200 µM L-ascorbic acid (Fisher Scientific BP351), 100 µM dibutyryl cAMP (Biogems 1698950), and 10 µM DAPT (Cayman 13197). After 35 days, organoids were transferred to ultra-low attachment plates (Corning 3516), with about 5 organoids per mL of media and cultured in DMEM/F12 with GlutaMAX supplemented with 1x N2, 1x B27 without Vitamin A, 10 ng/mL GDNF, 10 ng/mL BDNF, and 200 µM ascorbic acid.

Astrocytes were extracted from midbrain organoids starting at day 100. Organoids were gently dissociated in a trypsin enzyme solution (TrypLE Select, Gibco 12563011) using a glass pipette to break up organoids into large chunks. The organoid chunks from 5-10 organoids were placed on 15cm dishes coated with 0.1% gelatin and maintained in Astrocyte Medium (AM; ScienCell #1801). Astrocytes were allowed to grow out of the organoid chunks until the plate was confluent (about 1 week). At this point, astrocytes can be cryopreserved. Astrocytes were maintained in AM and switched to experimental media [1:1 DMEM/F-12:Neurobasal (Gibco 21103049), 1x B27 without Vitamin A, 1x N2, 1x MEM-Non-Essential Amino Acids (Gibco 11140050), 1x GlutaMAX (Gibco 35050061), 10 ng/mL CNTF (Peprotech 450-13)] 2-3 days before imaging.

### Astrocyte Viral Transduction

iPSC-derived astrocytes were transduced during regular passaging. In short, astrocytes were lifted, spun down, resuspended in AM media, and divided into 1.5mL microcentrifuge tubes, depending on the desired split ratio and number of transductions. The virus was added to the cell suspension and incubated for 5-10 minutes before plating the mixture. Media was topped off such that the final dilution of the virus in media was 1:50. Depending on the viral titer, this dilution sometimes increased or decreased. Astrocytes can also be transduced without passage by adding the virus directly to the media. Media is changed after 24 hours. Antibiotics were added to the media, if relevant, starting at least 3 days after transduction.

### MitoTracker and LysoTracker

For MitoTracker staining, astrocytes were incubated with 100 nM of MitoTracker DeepRed (Thermo Scientific M22425) for 30 min at 37 °C. For LysoTracker staining, astrocytes were incubated 100 nM of LysoTracker DeepRed (Thermo Scientific L12492) for 5 min at 37 °C. Cells were then washed twice with PBS before incubation with Hoechst 33342 (Thermo Scientific 62249).

### qPCR

RNA was extracted from samples using TRIzol (Invitrogen 15596018) and an RNA extraction kit following the manufacturer’s recommended protocol (Zymo R2062). RNA concentration and purity were measured on a spectrophotometer. cDNA was produced from 1 ug of RNA per sample, using Maxima H Minus Reverse Transcriptase (Thermo Scientific EP0752) and following the manufacturer’s recommendations. qPCR reactions were prepared in a 384-well plate (Applied Biosystems 4309849) with 1:10 dilution of cDNA, 2x PowerUp SYBR Green Master Mix (Applied Biosystems A25741) and 1 µM each of forward and reverse primers. qPCRs were run on QuantStudio 7 (Applied Biosystems 4485701) and analyzed using the ΔΔCt method, normalized to *ACTB* and *S18* rRNA. The primers can be found in the key resources table.

### Live Cell Imaging

In general, astrocytes were seeded at 10 x 10^3^ cells per well of a 96-well plate with a cover glass thickness polystyrene bottom (Greiner 655090) coated in 0.1% gelatin 3-5 days before imaging. Proliferation assays were seeded at 8 x 10^3^ cells per well and started 1 day after seeding. Before imaging, cells were incubated with Hoechst 33342 (Thermo Scientific 62249) or DRAQ5 (Thermo Scientific 52251) for 10 minutes.

Wide-field live cell imaging was performed using the Thermo Scientific CX7 High Content Screening Platform with 20x and 40x objective lenses (LUCPLFLN20x and LUCPLFLN40x). The microscope was equipped with an incubation unit that maintained temperature, CO_2_, and humidity during image acquisition. Confocal images were acquired on a Nikon Ti2E AX R confocal microscope with a 20x objective lens (PLAN APO λD 20x) and 6x optical zoom. Each well was imaged in 5 (confocal) or 25-36 (widefield) different areas, with the average of those images represented as a single data point.

### ImageJ Image Analysis

<u>Nuclear Proliferation:</u> Nuclear proliferation was quantified by converting images to a binary based on a threshold and then using ‘Analyze Particles’ to obtain a count for the number of nuclei in a field. The same threshold and parameters were kept for all time points.

<u>Mitochondrial Fragmentation:</u> Mitochondrial fragmentation was measured using the ImageJ plugin, Mitochondrial Analyzer ^104^. For thresholding, the “Block Size” was set to 1.45 microns and the “C-Value” was set to 5.

<u>Ratiometric Quantification:</u> Ratiometric quantifications for roGFP2, Timer, and Lemon were achieved by measuring the mean grey value in both channels of interest. The values for each channel were then divided to generate the ratio of relative intensity between the two channels.

<u>Kymograph Analysis:</u> Kymograph analysis of mitochondrial movement was achieved using the Image J plugins KymographClear 2.0 and KymographDirect ^105^. KymographClear 2.0 was used to trace mitochondrial trajectories and generate kymographs from which stationary and motile mitochondria were identified. KymographDirect was then used to extract quantitative information from the trajectories of the motile mitochondria.

### Cell Profiler Image Analysis ^72,106^

<u>Subcellular Co-localization:</u> For nuclear co-localization, nuclei in the genetically encoded fluorophore channel and the DRAQ5 channel were identified, and nuclei from cells that were not transduced were filtered out. For lysosomal and mitochondrial co-localization, propagation from the nuclear stain was used to identify cell bodies. Cells were filtered to only analyze those with lentivirus signal within the cell body. Co-localization was reported as the Mander’s coefficient indicating the overlap of the genetically encoded fluorophore with the commercially available stain.

<u>Lysosomal Distribution:</u> Cells were defined by nuclei and full cell fluorophore signal. Lamp1-mCherry signal was assigned to the cell body with which it overlapped. Only cells that were successfully transduced were included in the analysis. The distribution of the Lamp1-mCherry signal was calculated using the “MeasureObjectIntensityDistribution” with two bins generated for each cell using the nuclei as the center of the cell. Bins were scaled such that the inner bin (perinuclear) was always the same percentage of total cell area regardless of cell size.

### Quantification and statistical analysis

All experiments were conducted with 3-6 replicates. All quantifications were represented as mean ± standard deviation, except for the violin plots of the kymograph analysis. Details on data visualization, sample size, and images per replicate are included in figure captions. Details on statistical tests and significance are included in the text. For all statistical tests, *p < 0.05, **p < 0.01, ***p < 0.001, ****p < 0.0001. All statistical tests were conducted in *GraphPad Prism* and graphs were generated in *GraphPad Prism*.

## Supplemental Information

**Video S1: Mitochondrial axonal transport in *PRKN^+/+^*and *PRKN^-/-^* human iPSC derived neurons** Representative videos of COX8A-Emerald labeled mitochondria moving within neuronal axons over the course of 1 hour. Images were taken every 10 seconds and sped up 70x. Scale bar = 25 µm. Related to Figure 4.

---

**Fig. S1:**
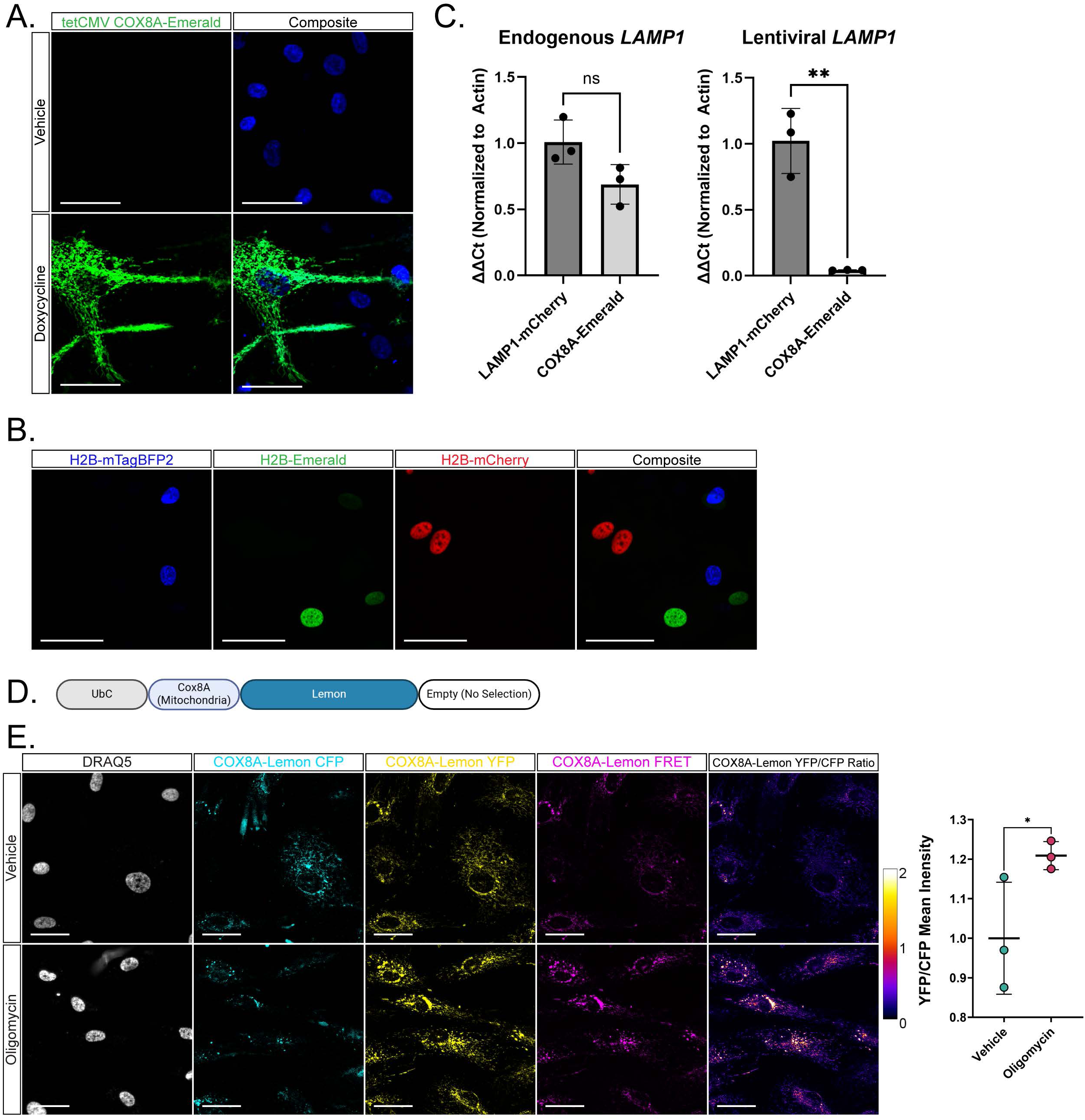
GEM-SCOPe supports a diversity of modifications and applications. **(A)** Representative images of astrocytes expressing COX8A-Emerald under the control of a doxycycline inducible promoter. Astrocytes were treated with a vehicle (DMSO; top) or doxycycline (bottom) for 72 hours. Nuclei were stained with Hoechst33342. **(B)** Representative images of a mixed population of astrocytes with nuclei expressing different H2B-fusion fluorophores. Astrocytes were transduced with a single lentivirus and then plated together. **(C)** qPCR for endogenous *LAMP1* (left) and lentiviral *LAMP1* (right) on RNA from astrocytes transduced with a virus localized to the lysosome (LAMP1-mCherry) or a virus not localized to the lysosome (COX8A-Emerald). Endogenous *LAMP1* qPCR primers were designed to amplify a sequence of *LAMP1* that is not included in the lentivirus and lentiviral *LAMP1* qPCR primers were designed to amplify the signal sequence of *LAMP1* and the beginning of the mCherry sequence. Bars represent mean and error bars represent standard deviation (n = 3 independent transductions). Statistical differences were determined by unpaired t-test; (left) p = 0.07; (right) p = 0.002. **(D)** Schematic of the lentiviral construct used in the following panels localizing Lemon, a pH responsive fluorophore, to the mitochondria with the targeting sequence of COX8A. Lemon undergoes Forster Resonance Energy Transfer (FRET) under alkaline conditions. **(E)** Representative images and quantification of astrocytes transduced with COX8A-Lemon and treated with a vehicle (DMSO; top) or 2 µM oligomycin (bottom) for 24 hours. Images were acquired with a 455nm-excitation laser and a 515-excitation laser. Far right: ratiometric representation of the emission in the yellow channel divided by the emission in the cyan channel and then pseudo colored so that orange, yellow and white indicate more relative yellow emission (more alkaline) while black and purple represent more cyan emission (more acidic). Nuclei were stained with DRAQ5. Central bars represent mean and error bars represent standard deviation (n = 3; each replicate is an average 36 images). Scale bars = 50 µm. For all graphs, * p< 0.05, ** p < 0.01, *** p < 0.001, **** < 0.0001.

**Fig. S2:**
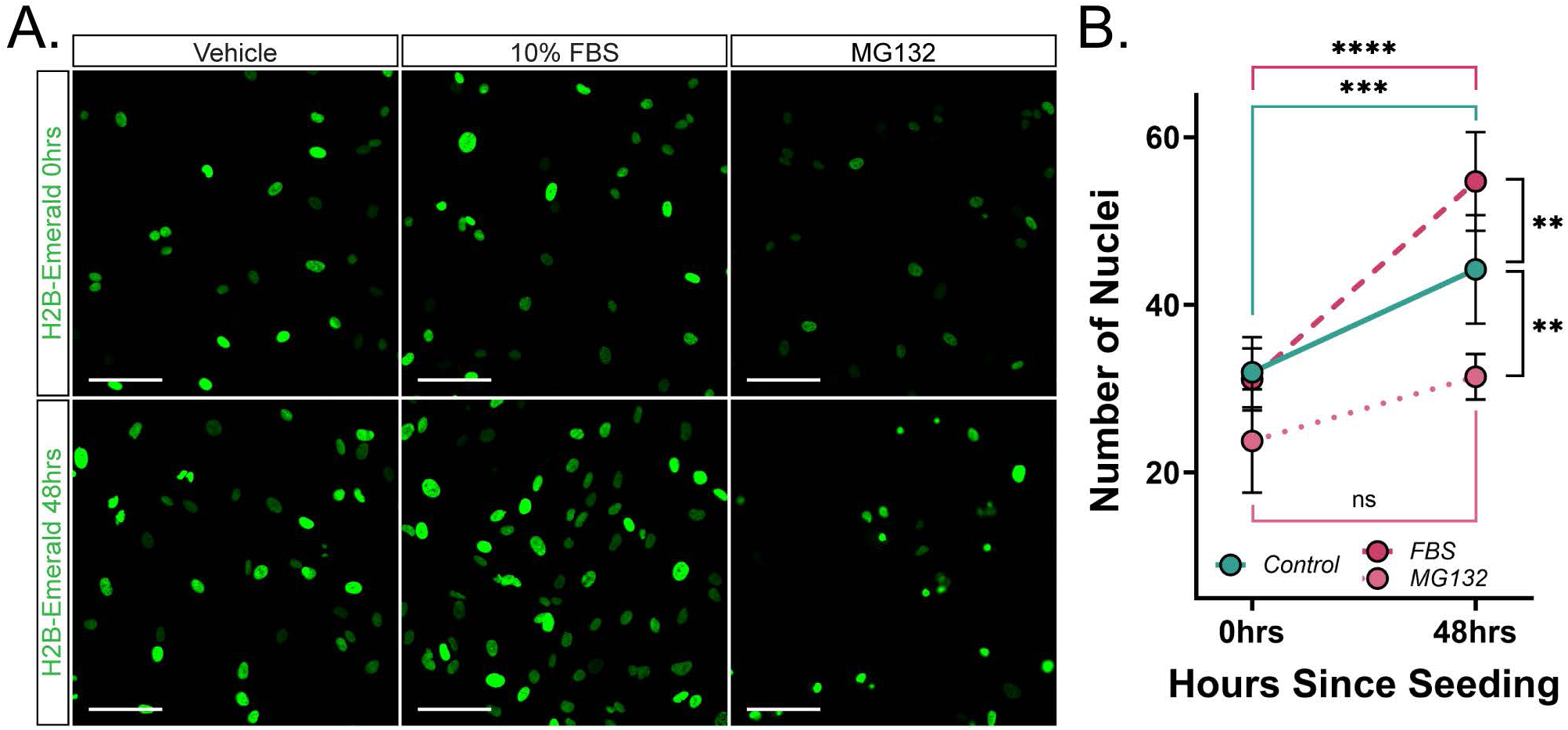
Astrocyte proliferation is activated by FBS and inhibited by MG-132. **(A)** Representative images of astrocytes transduced with H2B-Emerald lentivirus after 0 hrs (top) or 48 hrs (bottom) of treatment with a vehicle (DMSO; left), 10% fetal bovine serum (FBS; middle), or MG132 (right). Scale bars = 50 µm. **(B)** Quantification of number of H2B-Emerald nuclei per image field 0 hours and 48 hours after treatment. Dots represent mean values and error bars represent standard deviation (n = 6 (n = 3 for MG-132 treatment) per time point; each replicate is an average over 25 images). Scale bars = 50 µm. All images were acquired on the CX7 HCS platform with a 20x objective lens. For all graphs, * p< 0.05, ** p < 0.01, *** p < 0.001, **** < 0.0001.

**Fig. S3:**
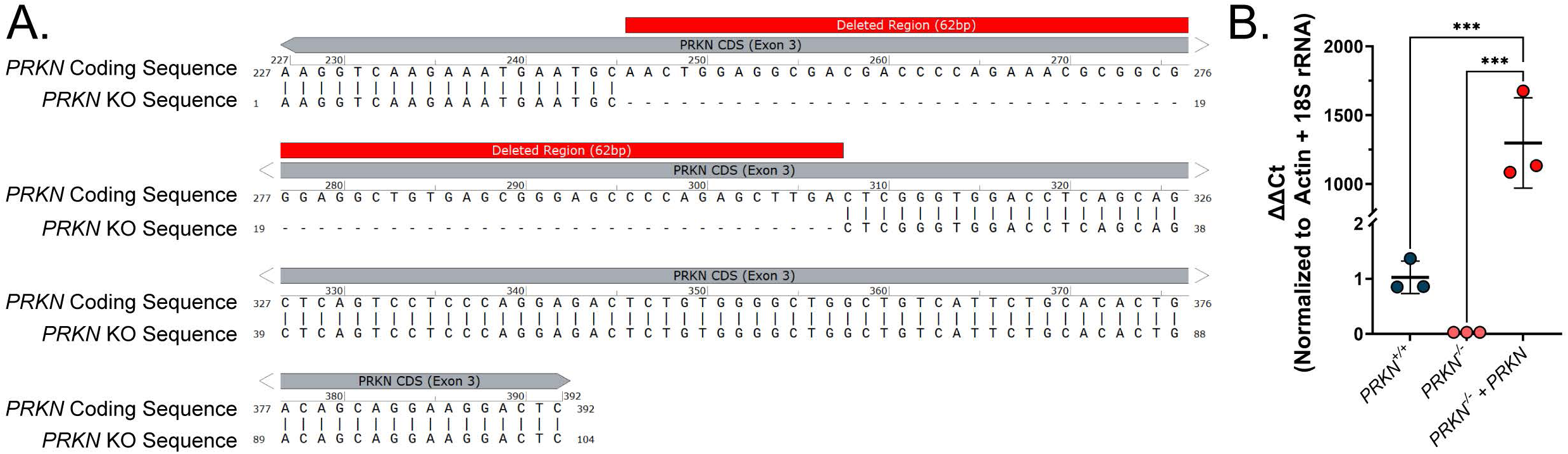
***PRKN* knockout and overexpression (A)** Sequencing alignment of *PRKN*^-/-^ astrocytes with the *PRKN* coding sequence showing a 62bp deletion in exon 3 **(B)** qPCR for *PRKN* in *PRKN^+/+^, PRKN^-/-^*, and *PRKN^-/-^* with *PRKN-*overexpression. qPCR primers were designed to amplify the region deleted in the *PRKN^-/-^* cells. Bars represent mean and error bars represent standard deviation (1-way ANOVA with Tukey’s HSD; *PRKN^+/+^* vs *PRKN* knock-in: p = 0.0004; *PRKN^-/-^* vs *PRKN* knock-in: p = 0.0004; n = 3 independent transductions). For all graphs, * p< 0.05, ** p < 0.01, *** p < 0.001, **** < 0.0001.

**Fig. S4:**
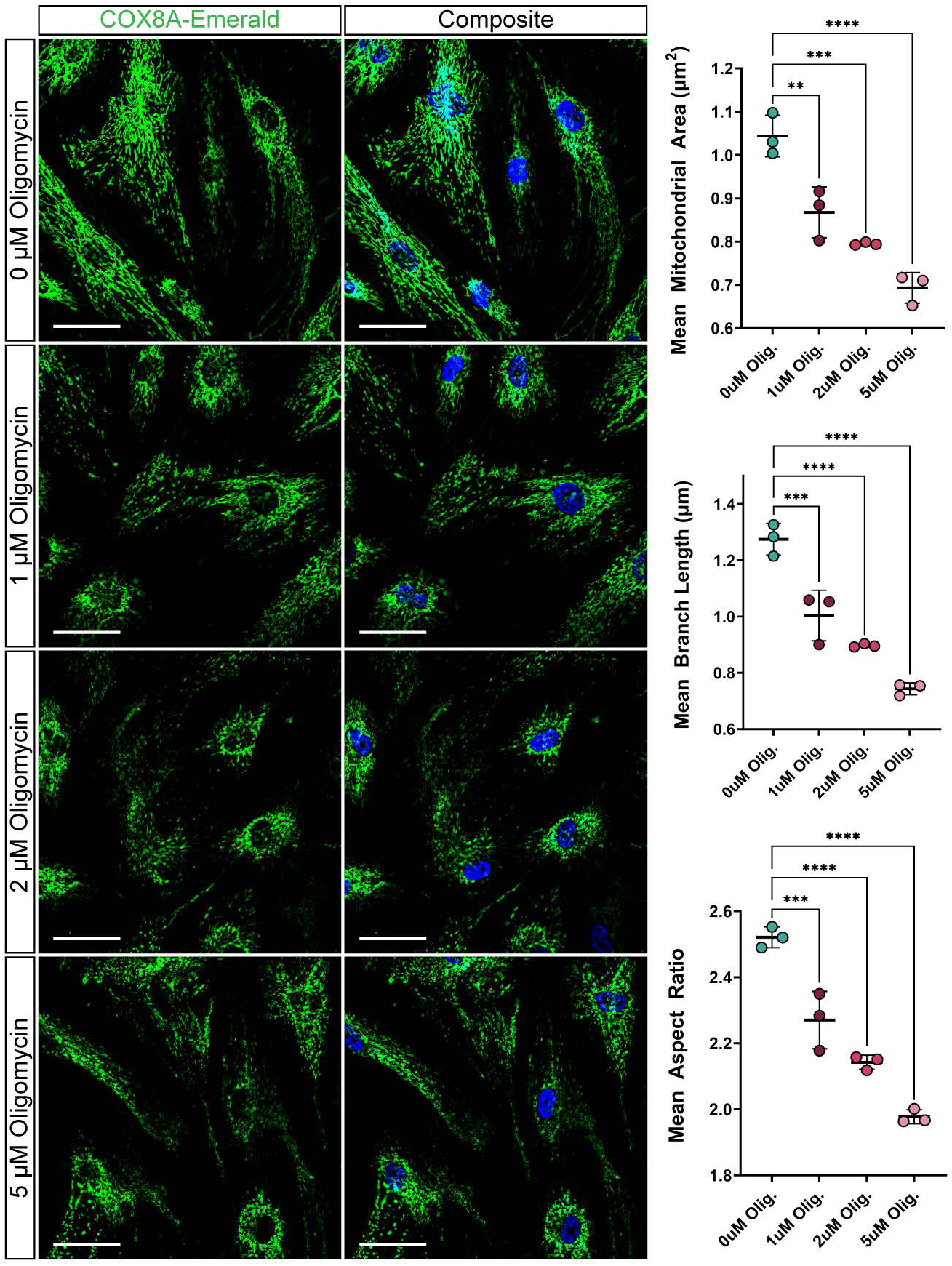
Mitochondrial fragmentation exhibits a dose dependent response to oligomycin-induced stress. **(A)** Representative images of astrocytes transduced with COX8A-Emerald and treated with a vehicle (DMSO), 1µM, 2 µM, or 5 µM oligomycin for 4 hours. Nuclei were stained with Hoechst33342. Scale bars = 50 µm **(B)** Quantification of mitochondrial features indicative of mitochondrial network fragmentation: mean area, perimeter, aspect ratio, number of branches, branch length, and number of branch endpoints. Central bars represent mean and error bars represent standard deviation (1-Way ANOVAs with Dunnett’s test; n = 3; each replicate is an average over 36 images). For all graphs, * p< 0.05, ** p < 0.01, *** p < 0.001, **** < 0.0001.

**Table S1:** A list of all plasmids generated in this study. . Details for all plasmids, including subcellular localization, fluorophore, antibiotic resistance, and in which figure each plasmid was used. More plasmids were generated as part of GEM-SCOPe than used in the final publication, and these are also included here and available on Addgene.

| Plasmid Name | AddGene ID | Localization | Fluorophore | Antibiotic Resistance | Relevant Figures |
| --- | --- | --- | --- | --- | --- |
| pFUW H2B-mTagBFP2-p2A-PuroR | 239552 | Nucleus | mTagBFP2 | Puromycin | 1, 2, S1 |
| pFUW H2B-Emerald-p2A-PuroR | 239553 | Nucleus | Emerald | Puromycin | 2, S1, S2 |
| pFUW H2B-mCherry-p2A-PuroR | 239554 | Nucleus | mCherry | Puromycin | 2, S1 |
| pFUW LAMP1Sig-Emerald | 239555 | Lysosome | Emerald | NA | 1, 3 |
| pFUW LAMP1Sig-mCherry | 239556 | Lysosome | mCherry | NA | 1 |
| pFUW COX8ASig-Emerald | 239557 | Mitochondria | Emerald | NA | 1, 4, S4 |
| pFUW COX8ASig-mCherry | 239558 | Mitochondria | mCherry | NA | 1 |
| pFUW PuroR-P2A-COX8ASig-Timer | 239559 | Mitochondria | Timer | Puromycin | 5 |
| pFUW COX8ASig-Lemon | 239560 | Mitochondria | Lemon | NA | S1 |
| pFUW mito-GRX1-roGFP2 | 239561 | Mitochondria | roGFP2 | NA | 6 |
| pFUW mTagBFP2 | 239562 | NA | mTagBFP2 | NA | NA |
| pFUW Emerald | 239563 | NA | Emerald | NA | NA |
| pFUW mCherry | 239564 | NA | mCherry | NA | NA |
| pFUW mTagBFP2-P2A-PuroR | 239565 | NA | mTagBFP2 | Puromycin | NA |
| pFUW Emerald-P2A-PuroR | 239566 | NA | Emerald | Puromycin | 3 |
| pFUW mCherry-P2A-PuroR | 239567 | NA | mCherry | Puromycin | NA |
| pFUW mTagBFP2-P2A-NeoR | 239568 | NA | mTagBFP2 | Neomycin | NA |
| pFUW Emerald-P2A-NeoR | 239569 | NA | Emerald | Neomycin | NA |
| pFUW mCherry-P2A-NeoR | 239570 | NA | mCherry | Neomycin | NA |
| pFUW PRKN-P2A-PuroR | 239571 | NA | NA | Puromycin | 6, S3 |

